# A Scalable Data Access Layer to Manage Structured Heterogeneous Biomedical Data

**DOI:** 10.1101/067371

**Authors:** Giovanni Delussu, Luca Lianas, Francesca Frexia, Gianluigi Zanetti

## Abstract

This work presents a scalable data access layer, called PyEHR, intended for building data management systems for secondary use of structured heterogeneous biomedical and clinical data. PyEHR adopts openEHR formalisms to guarantee the decoupling of data descriptions from implementation details and exploits structures indexing to speed up searches. The persistence is guarantee by a driver layer with a common driver interface. Presently, are implemented the interfaces with two NoSQL DBMS: MongoDB and Elasticsearch. The scalability of PyEHR has been evaluated experimentally through two types of tests, namely constant load and constant number of records, with queries of increasing complexity on a two synthetic datasets of ten millions records each, containing very complex openEHR archetype structures, distributed on up to ten working nodes.

## I. INTRODUCTION

Next-generation sequencing(NGS) and other high throughput technologies are rapidly transforming life sciences [1], [2]. Their use is now routine in biology labs and is very quickly expanding to clinical research [3] and applications [4]. This transformation has led to an abrupt transition towards a data-intensive domain where sophisticated computational analysis is essential to extract biologically relevant information from the raw data [5], [6].Consequently, a great effort has been made to develop scalable computational tools able to cope with the current data load and the foreseen, much larger, one that is expected to arise due to the increasing use of these technologies in large-scale clinical studies [7], [8]. A specific aspect of these computations is that it is important to maintain full processing traces as an integral part of the results, since the latter is at the end of complex and deep computational pipelines whose outcome depends strongly on chosen parameters and configurations. This is particularly true now for the analysis of data coming from NGS and it will become equally relevant with the expected diffusion of Computer Aided Diagnosis systems [9]. Analogously, the explosive diffusion of digital data acquisition devices for biomedical applications, ranging from fully traced clinical procedures [10] to IOT personal health acquisition devices [11] has dramatically increased the amount of context information that can be attached to phenotypic information. In more general terms, all computed and phenotypic information typically map to deeply structured records with potentially recurring substructures, e.g., see figure 3. The development of novel algorithms, data and meta–data handling techniques needs, therefore, to be complemented by robust, scalable, computable, uniform and implementation-independent descriptions of deeply structured data [3]. Here, scalability is meant both with respect to the sheer size of the data and to the evolution of their structure and type. The latter issue is typical of longitudinal biomedical studies, where multiple, heterogeneous data types may become necessary within the time span of the project. Ideally, common solutions based on ad-hoc database tables should be replaced by computable formalisms for the meta-description of structured records. Such systems should easily support operations such as aggregations on sophisticated profile descriptions across all the available records, as well as in-depth navigation of all data related to a specific study participant.

In this paper, we describe PyEHR, a data access layer for the creation of data-management systems for biomedical research specifically designed to efficiently manage queries on large collections of very heterogeneous structured records with recurring repeated substructures. The system is designed to be scalable with respect to the evolution and heterogeneity of biomedical research data structures and to data volumes compatible with regional-scale studies. Our motivation for the development of PyEHR comes from our direct experience in providing computational support to a wide range of biomedical research projects, including large-scale genome sequencing studies [12]–[14] and safety assessments of novel gene therapy approaches [15] as well as the running of a large scale NGS facility [16].

We use openEHR [17], a computable formalism for the meta-description of structured clinical records, as a systematic approach for handling data heterogeneity and to have a clear separation between a well defined semantic description of data structures and the actual storage implementation. Scalability with respect to dataset size, on the other hand, is achieved through a multi-tier architecture with interfaces for multiple data storage systems. With this approach, it is possible to express queries at the semantic level and to perform structural optimizations well before they are translated to a specific implementation. Currently, the specific DBMS supported are two NoSQL databases/search engines, MongoDB [18] and ElasticSearch [19], but other backends, even relational ones, are easily implementable.

We are interested in traversing data, in a heterogeneous but structured data lake [20], efficiently and in a scalable manner, trying to exploit as much as possible the structure of the queried data. Our approach is based, in practice, on indexing each unique class of structures to speed-up the search process and decouple as much as possible the logical description of the operation that should be performed from the specific data engine that lies beneath. The specific context in which we consider the problem is, of course, biomedical research but the discussion is, in itself, quite general.

PyEHR has been tested on a set of synthetic data, created to challenge it with very deep and complex structures. As a reference, we performed the same query tests using a straightforward search approach implemented as an Apache Hadoop Mapreduce application. Our results indicate that PyEHR has a good scalability.

The remainder of the paper is organized as follows.In section II, we provide a short discussion on the context. Section III, offers a brief description of the openEHR standard, its way of defining and querying data through the ADL and AQL languages and how it can represent complex data structures. In section IV is discussed how the indexing of structures has been inplemented in our system and what are its input and output. Section V is dedicated to a short presentation of the whole system, PyEHR. In the following section VI the scalability appraisal of PyEHR is addressed, from the data creation to the choice of tests and ultimately to their results. Next section VII compares and discusses the results. Then in section VIII is described the relevant related work found in literature. Finally section IX present our conclusions and suggested future foreseen activities whereas section X highlights what we reckon are the major limitations of this work.

## II. Context

Biomedical research and its clinical applications are quickly becoming data intensive enterprises [21]. On one hand the transition is fueled by the increased capability of generating big digital data. This is mainly due to technology innovations, increased adoption of electronic health records(EHR) and the creation and capture of, novel, patient generated data like those deriving from body sensor data, smartphone apps, social networks’ posts, etc.. Technology innovations, in particular, have enabled the automatic, and progressively fast and cheap, creation of big volumes of omics data and high resolution medical imaging. On the other hand there have been technological advances in storage and computing capabilities too, that allow to collect, save and analyze increasingly large amounts of data.

Often the biomedically relevant information is obtained as the end result of complex and deep computational pipelines that are in continuous evolution and whose outcomes depends strongly on chosen parameters, configurations, software release version, etc. [22]. It’s important to point out that the metadata defining the pipeline are as important as the results themselves; they are critical to understand for instance the reasons behind different outcomes or get test reproducibility.

Its importance is exemplified in figure 1 [23], which highlights the degree of dissimilarity between sets of genomic variants identified from the exome sequencing of parent-child trios [24]. All sets of variants were extracted from the same data using the same conceptual analysis protocol with different versions of the same software tools and/or reference genomes. The pattern shown is typical of data-intensive analysis pipelines: the evolution of analysis algorithms and reference datasets has a drastic impact on results. Therefore, to effectively use the data it is necessary to be able to recalculate results with the new best available tools and models, which in turn implies having full provenance information.

**Fig. 1.**
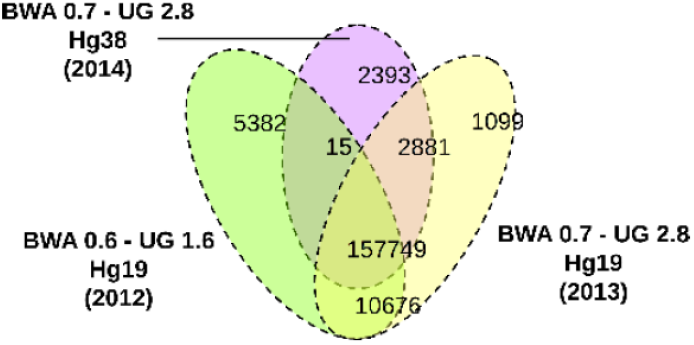
Number of common variants produces by different pipelines given identical input [23].

The result itself of the genomic variants identification is a deeply structured dataset see figure 2. It is important to note how the data structure uses, e.g., to annotate variants, recurrent substructures, highlighted in figure with a dash line.

**Fig. 2.**
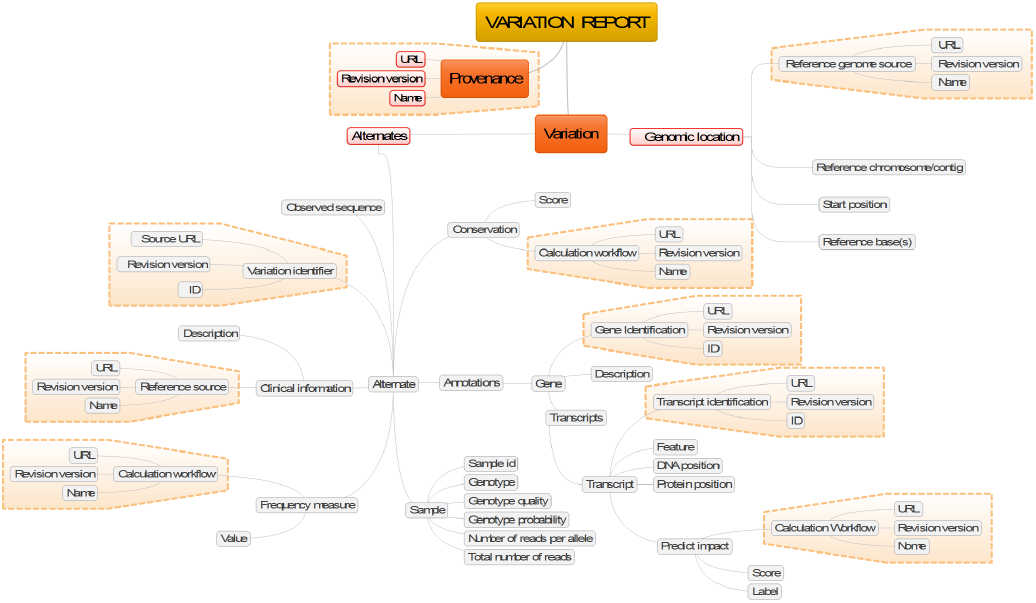
Mind map of a set of annotations produced by a variant calling procedure.

At the same time clinical information, too, is becoming more and more structured. Figure 3 shows an example, related to digital pathology [25], of a complex chain of dependencies that should be fully maintained, and accurately described, to precisely relate the actual data, here a whole slide digital image, to the relevant context. A parent specimen extracted from a patient can have multiple specimens children divided into blocks which are sliced. The resulting slides can have one item per container, multiple items from the same block or items from different parts in the same block.

**Fig. 3.**
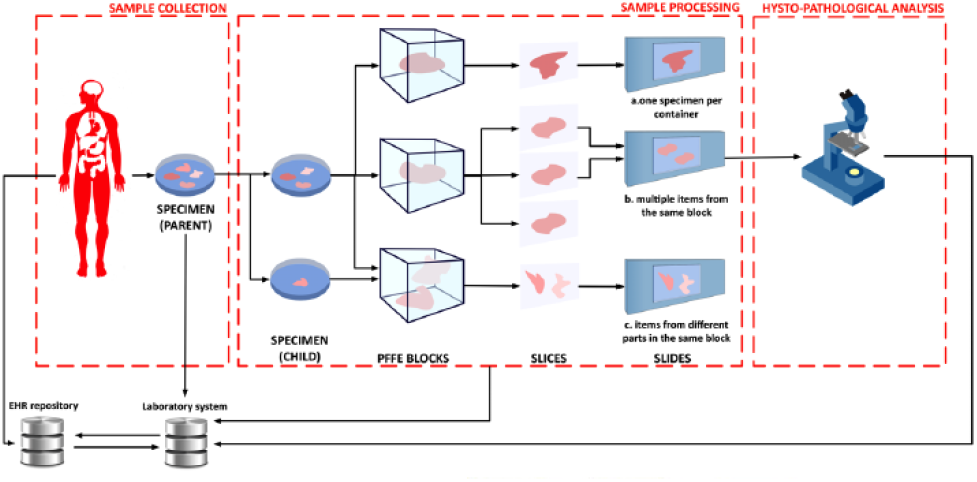
Example of slides acquisition in digital pathology.

In practice, the gathering of phenotypic information, acquired raw data and analysis results, produces the accumulation of vast quantities of heterogeneous structured (e.g., reports, analysis provenance graphs) and non-structured data (e.g., raw data, images) [26]–[28]. Given the continuous evolution of data sources, it is not technically feasible to map the problem to a data-warehouse hence data lakes could be used instead [22], [29]. The major advantages of data lakes over the standard data-warehouses are their flexibility, as they can contain any type of data like structured, unstructured or raw, their robustness, because they only require a schema-on-read and are easily reconfigured, and their easy and affordable scalability [30].

## III. OpenEHR

OpenEHR is a standard evolved from the Australian Good Electronic Health Record(GEHR), whose most peculiar feature is the two-level framework, where the information model or reference model (RM) is kept separated from the clinical knowledge or Archetype Model (AM), see figure 4. The information model defines the generic types and structures for record management. It is designed to be limited to domain-invariant data elements and structures, such as quantity, coded text and various generic containment structures, the classes. The AM provides, through archetypes and templates, structures and admissible values of a series of fields with well-defined semantic relationships. The archetypes are expressed using constraints on instances of the underlying reference model while the templates are models of content corresponding to use-case specific data sets, constituted from archetype elements. The separation of the two model has the advantage of separating responsibilities in knowledge management and sharing. Medical professionals can author archetypes and IT programmers can develop the storage and sharing layer. The architecture is designed to make use, optionally, of external health terminologies, such as SNOMED CT, LOINC and ICDx. Each archetype is a computable definition, or specification, for a single, discrete clinical concept and is expressed in Archetype Definition Language (ADL) [31], which is an ISO standard, but able to be viewed and reviewed in ‘clinician-friendly’ formats, as structured definitions and mind maps. They represent some real world concept, such as ‘‘patient”, “blood pressure”, or “antenatal examination”. Archetyped data have the same meaning no matter what context is used within the EHR and, similarly, no matter which EHR system is employed or what language is adopted. ADL can be used to write archetypes for any domain where formal object models exist which describe data instances. Archetypes are language-neutral, and can be authored in and translated into any language. ADL encloses three other syntaxes, cADL (constraint form of ADL), dADL (data definition form of ADL), and a version of first-order predicate logic (FOPL), to describe constraints on data which are instances of some information model. In figure 5 is shown an example of a fabricated archetype. In the example the idea of a sport racket is defined in terms of constraints on a generic model of the concept DEVICE. All sections are written in dADL except the “definition” that adopts the cADL. In ADL the main sections are called concept”, “language”, “definition” and “ontology”. If the racket had been derived from another archetype then a “specialize” section would have been added, something like the inheritance in programming languages; a new archetype derived from that of the parent by adding a new section to its domain concept section. The “definition” contains the main formal definition of the archetype. Inside, referring to the example, there is a class (“DEVICE”) and its attributes (“length”,”width”, etc.), created in the object model associated to this archetype. The attribute part has a constraint called cardinality, applicable to any attribute, that indicates limits on the number of members of instances of container types such as lists and sets. In the example are allowed from zero to many instances of parts. Another important constraint, missing in the example, is the “ARCHETYPE_SLOT” that admits one or a list of archetypes to be included within an archetype definition. That means that archetypes can be nested and combined to describe complex structures like the one we are facing when dealing with heuristic biomedical pipelines. At last there is the “ontology” section, which explains what the local nodes, elements that conventionally starts with AT, and the local constraints, that conventionally starts with AC, mean and how they are bound. In the latest release of ADL, 2.0, this section has been renamed “terminology”. It can be noted the repeated object/attribute hierarchical structure of an archetype that provides the basis for using paths to reference any node in an archetype, paths that follow a syntax subset of W3C Xpath.

**Fig. 4.**
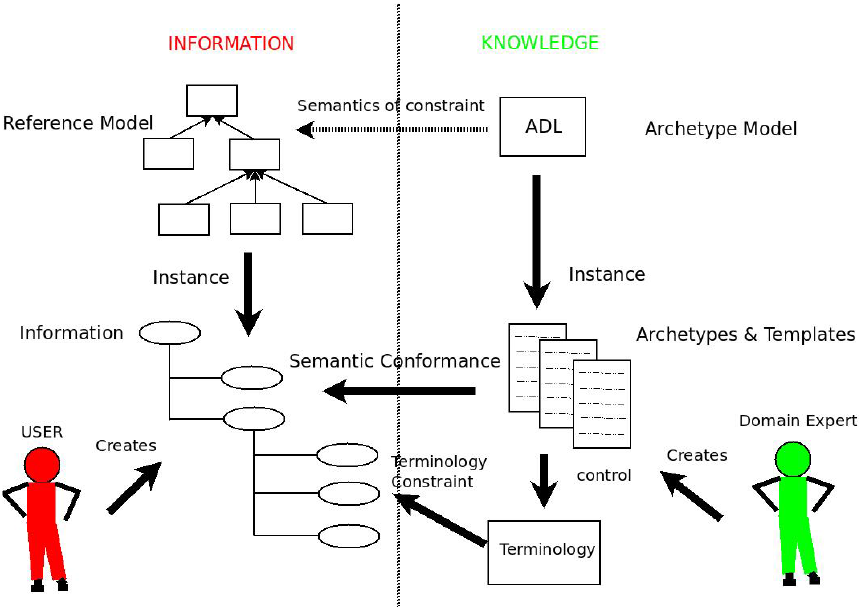
The openEHR two-level separation

**Fig. 5.**
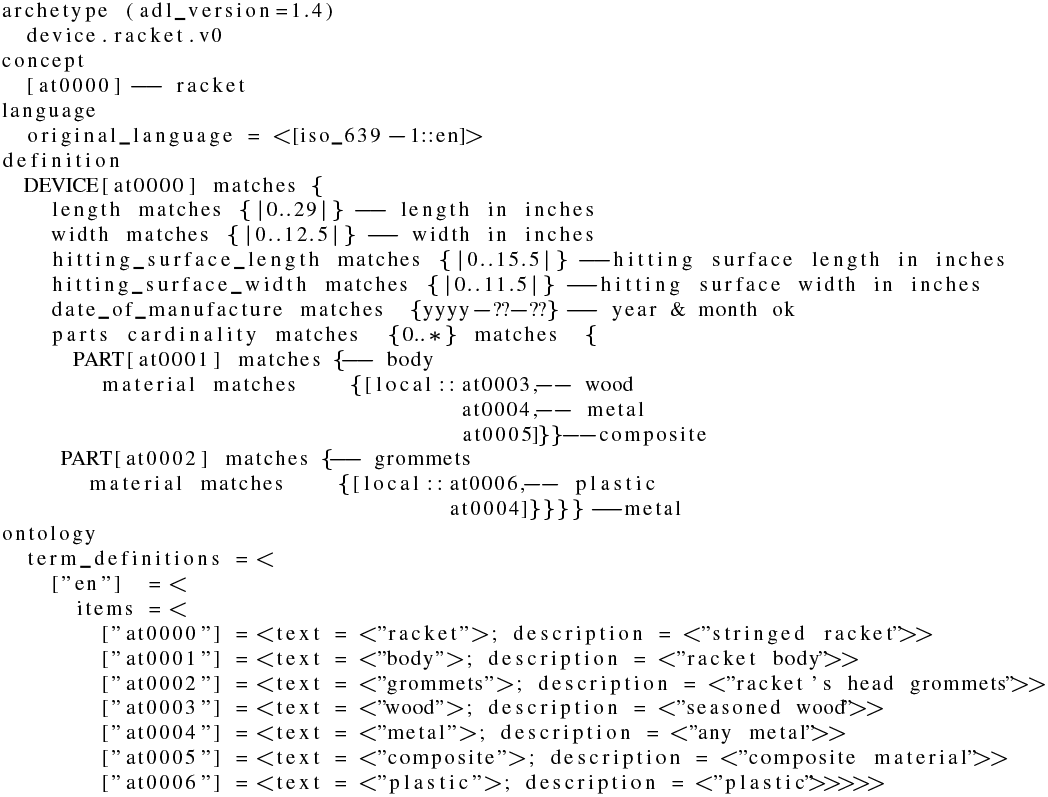
Example of an archetype expressed in ADL.

The data expressed as single archetypes or more frequently compositions of archetypes, can be queried in AQL [32], the archetype query language. AQL, formerly known as EHR Query Language (EQL), is a declarative query language born in 2005 to answer to the following four requirements for an archetype-based query language [33]:

1) the query language should be able to express queries for requesting any data item from an archetype-based system, i.e. data defined in archetypes and/or the underlying reference model;
2) the query language should be able to be used by both domain professionals and software developers;
3) the query language should be portable, i.e. neutral to system implementation, application environment and programming language;
4) the syntax should be neutral with respect to the reference model, i.e. the common data model of the information being queried. Particular queries will of course be specific to a reference model.

The main asset of AQL is that, unlike other query languages such as SQL or XQuery, it allows to express the queries at the archetype level, i.e., semantic level, other than at the data instance level. This is the key to easily share queries across system boundaries or enterprise boundaries. Its main features are:

- the utilization of openEHR archetype path syntax in the query expression;
- the utilization of containment mechanisms to indicate the data hierarchy;
- the utilization of ADL-like operator syntax, such as matches, exists, in, negation;
- a neutral expression syntax. AQL does not have any dependencies on the underlying RM (Reference Model) of the archetypes. It is neutral to system implementation and environment;
- the support of queries with logical time-based data rollback.

AQL has clauses similar to the well-known SQL, and in particular they are: SELECT, FROM, WHERE, ORDER BY, TIMEWINDOW. The SELECT clause specifies the data elements to be returned, using openEHR path syntax to indicate expected archetypes, elements, or data values. The FROM clause specifies the data source and the containment constraints introduced by the CONTAINS keyword. The WHERE clause defines, within the chosen source, data value criteria. The ORDER BY clause is used to select the data items that rule the returned result set order. Finally, the TIMEWINDOW clause restricts the query to the specified time or interval. An example of an AQL query is given in figure 6. The domain of sport matches “Matches” is searched for the ones where was used a wood racket whose length exceeds twenty-eight inches and if they exist the query returns their match identifier. The CONTAINS keywords allow to navigate through the nested archetype. In particular in the example the archetype racket must be inside a specified OBSERVATION archetype (test.OBSERVATION.devices.v23) which in turn must be inside a specified COMPOSITION archetype (test.COMPOSITION.encounter.v23). Between the two there may be any number of archetypes or no archetypes at all.

**Fig. 6.**
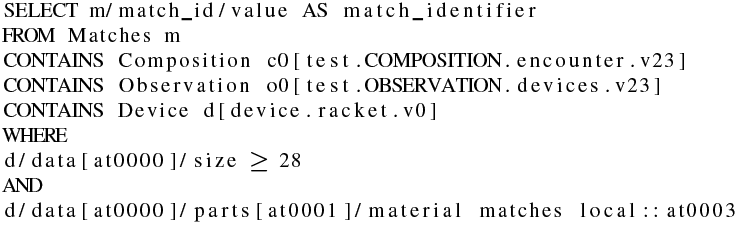
Example of an archetype query in AQL.

## IV. Data Structures Indexing

OpenEHR provides a valid way of retain data semantics, context and provenance but something has to be done in order to search this data efficiently and in a scalable way. Given the complex structure of the data, deep, nested and rich of repetition blocks, we thought to implement in our system an indexing of, unique, structures to greatly ease the task of the db engine during queries.

For example figure 7 shows an archetype that represent a certain class of structures. In the structure each node, denoted by given shape and capital letter, is again an archetype. Different shapes/letters mean different archetypes while the text “0..*” implies that is possible to have from none to any number of the following archetypes. In figure 8 are displayed some of the structures instances of the archetype of figure 7. The number inside the shaped node, next to the capital letter, indicates a given instantiation. We have instances with no octagone nodes and instances with one or two archetypes of that type. This is very difficult to be mapped in a traditional relational dbms as its scheme changes hence any implementation would be temporary and fragile. Whatever type of dbms is used indexing the structures saves time while traversing the data tree looking for query matches. In practice, in order to get our structures index, each data structure is pruned of non–structural details, analyzed for the mutual positions of archetypes, and transformed to a canonical description. The latter are given an identifier and stored in a specialized database. This is one of the responsibilities of the Index Service. Archetypes that occupy different places in a list of items, that is permutations, are assigned the same structure id, avoiding an overload of structures that represent in fact the same arrangement. In figure 9 is shown an example of a structure that contains six archetypes differently nested inside the containing archetype openEHR-EHR-COMPOSITION.encounter.v1.lbl-00001. The “class” property defines the archetype name whereas the “path_from_parent” property indicates a path relative to the closest containing archetype. It can be noted that in the example all archetypes, except the root archetype, have the same “path_from_parent” property and the reason is that they refer to the same type of archetype, a openEHR-EHR-COMPOSITION.encounter, whose path to the contained archetype is always the same. During the query process the Index Service is asked to perform another task. Let’s suppose to have an AQL query like the one shown in figure 10 and to apply that to the previously defined structure of figure 9. The Index Service look for all the structures that match the containment constraints and returns a list of their ids together with the absolute path or paths to get to the innermost archetype of the CONTAINS clause. Putting it to use in the example, the structure of figure 9 matches the containment constraint so are returned the unique id of the structure and the path, only one here, obtained concatenating all the archetypes’ relative paths from the root to the last contained archetypes. The Query Management System thus has all the information to search for the data that comply to the WHERE clause examining only a subset of all the stored records, i.e. the ones whose structure satisfies the containment clauses, and knowing in advance where to look for, return more quickly the data chosen in the SELECT clause.

**Fig. 7.**
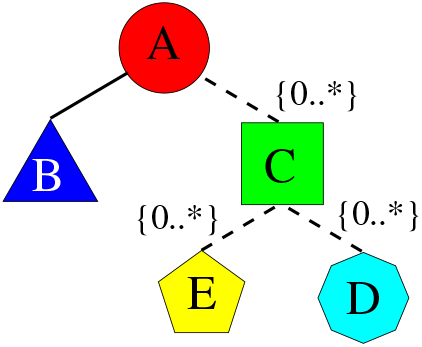
Example of an archetype, composed of other archetypes, shown as a tree structure.

**Fig. 7.**
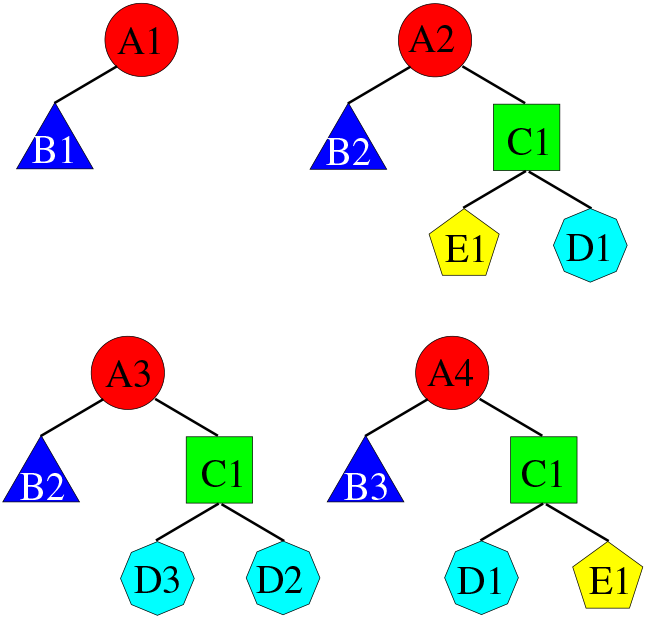
Some possible instances of the archetype of figure 7.

**Fig. 9.**
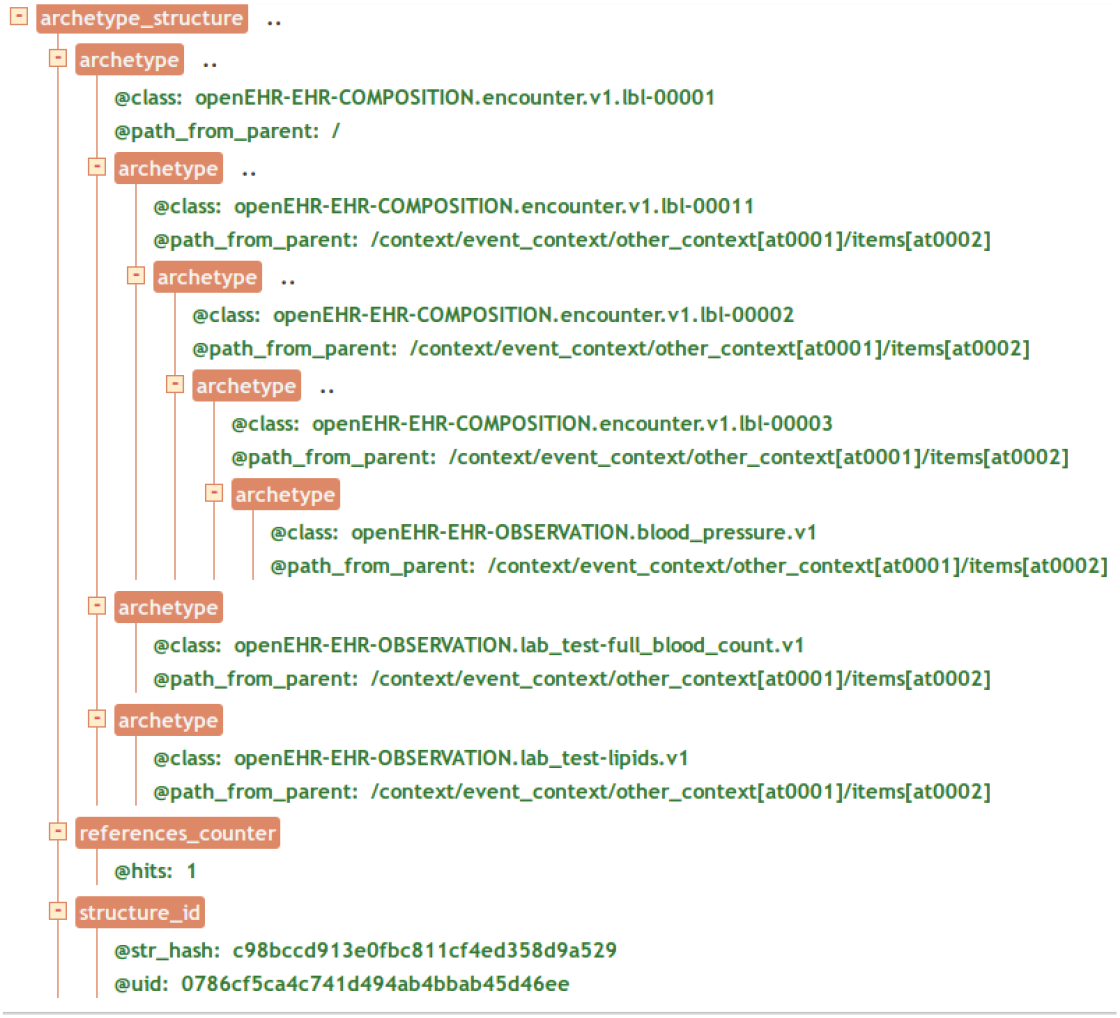
Tree view of a data structure. (See section X for credit)

**Fig. 10.**
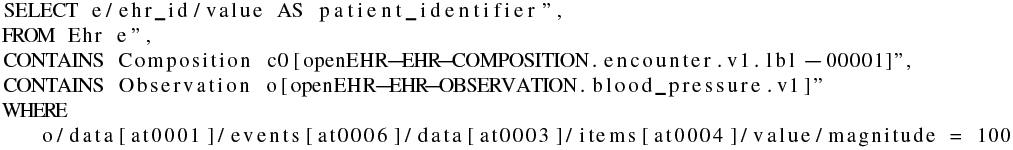
AQL query for the data structure in figure 9

We found that our structures indexing has a low impact on insertion while provides a big boost in querying performances.

## V. Implementation

PyEHR, in a nutshell, is an open source data access layer designed to help building applications for secondary use of clinical and biomedical data. Nevertheless it can also be used as it is. The software has been written mostly in Python, with services exported through a REST architecture. The target, as we already mentioned, is to be able to treat complex heterogeneous structured data. The system implements the openEHR standard, storing archetypes representations and accepting queries in AQL. This gives flexibility both in accepting and searching complex data structures whereas indexing speeds up the process of querying big amount of data. PyEHR make use of an external database, named Ehr Database, to permanently store the clinical and biomedical data. The driver interface has been conceived to easily allow the deployment of any database, an idea already adopted in [34]. In fact a new database driver needs only an entry in the driver’s factory class (see listing in figure 11) and the implementation of a short list of routines according to a common driver application program interface (API). Currently we are supporting two NoSQL database management systems: MongoDB [18] and ElasticSearch [19]. Both systems, having been designed to handle hierarchical sets of key-value items, are easily adaptable to the document-like structures of openEHR data. In figure 12 are shown the main tasks accomplished by the driver API. For clarity are omitted all the functions related to versioning control. Each driver has the following responsibilities:

- manage connections and disconnections to the Ehr Database;
- provide full CRUD (Create, Read, Update, Delete) support;
- handle queries (rebuilding them in the driver natural query language and executing them);
- encode/decode data to/from the wrapper objects defined in the services layer, automatically converting any special characters;
- create data structures such as SQL tables or folders;
- split or join records as required by the underlying storage system.

**Fig. 11.**
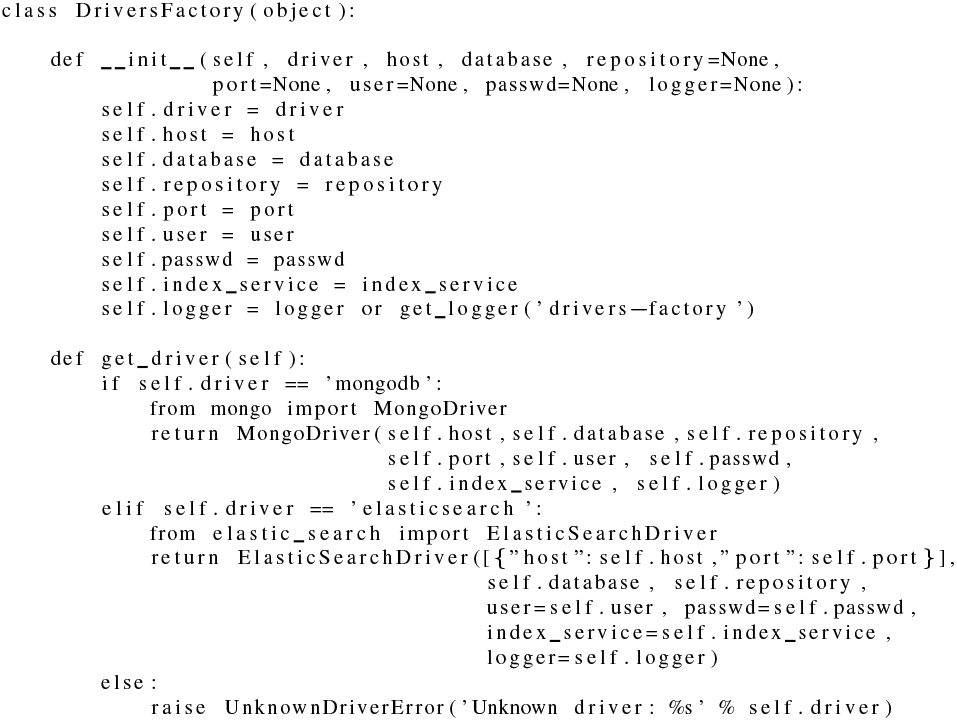
Drivers Factory Class

**Fig. 12.**
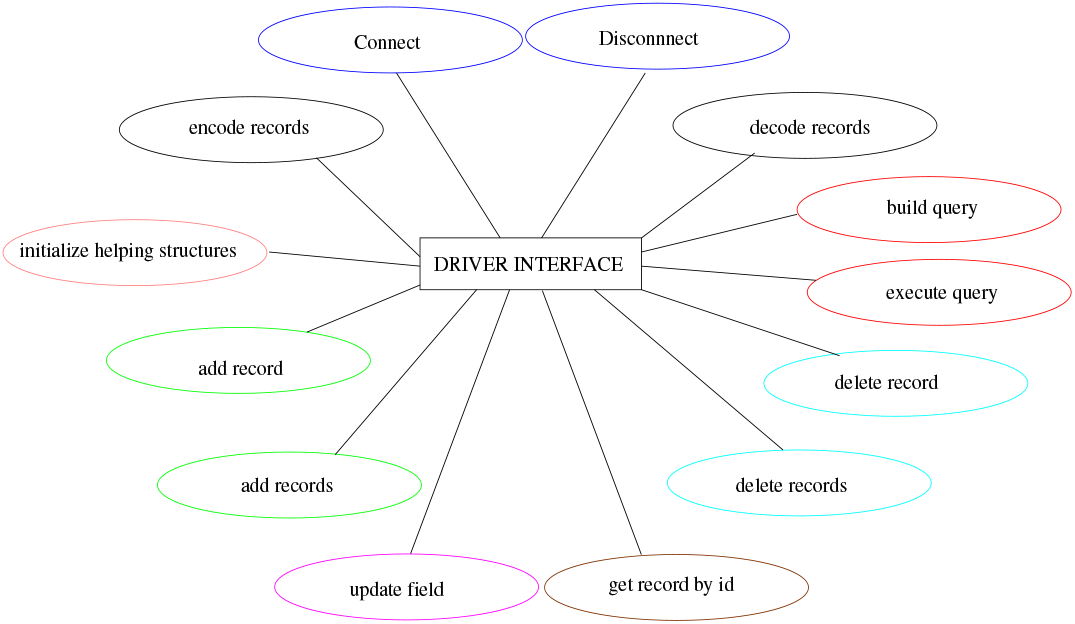
Main driver API tasks.

Sometimes the driver interface must implement solutions to overcome database management system limitations or undesired features. For instance Elasticsearch does not allow that under the same “index”-”type” combination exist documents with the same name but different class, e.g., foo defined as an integer variable in a document and as a dictionary in another. In order to overcome what was, for our purposes, a limitation we had to change the type name, creating a unique one by associating a generic name to the structure id, but this in turn has required the creation of a lookup table for quickly retrieving the “index”-”type” couple information for a given document id, an event very frequent especially while managing documents’ versioning. The service layer has its own representation of data objects so the driver, in order to manage them acts, also, as a translating interface that converts to and from the well-known standard json. This allows the data objects to be handled by the REST API and the db engine though their data representation may differ. An example are clinical records that are represented internally as documents, whose structure closely match the original, written in ADL and are converted, when needed, to json representation and given in this format to the database driver or the REST API. PyEHR overall architecture is summarized in figure 13. There are three modules (Data Management System, Query Management System, Index Service) that interacts with each other and two databases (Ehr Database, Structures Database). The Structures Database, mentioned in section IV, stores unique data structures. The Ehr Database, meant to be interchangeable, stores the biomedical data along with other metadata, including an id representing their structure that links data and structures databases. The Index Service, described in section IV, has been implemented using BaseX [35] as database for managing and querying index structures.

**Fig. 13.**
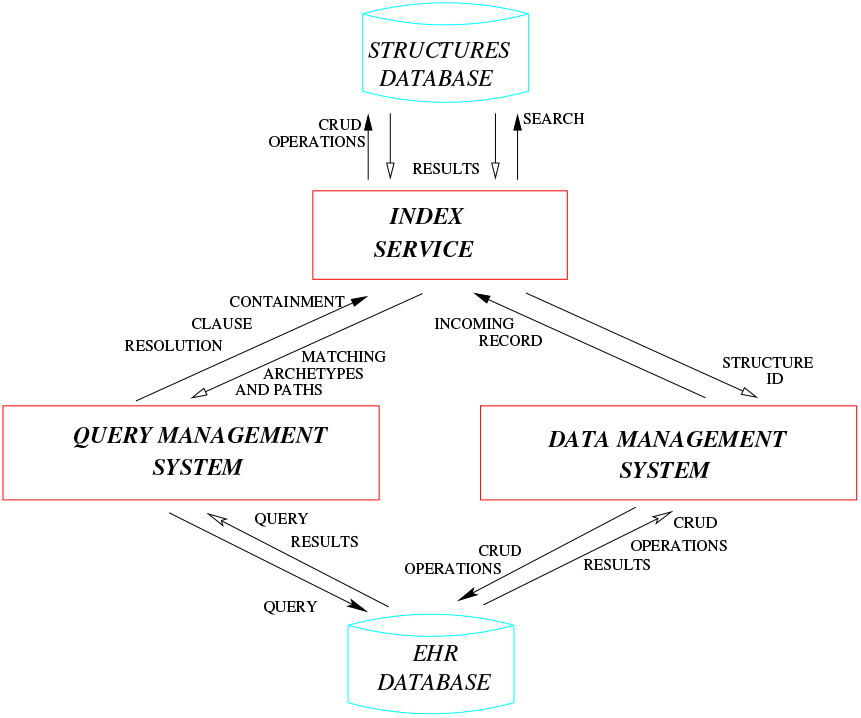
PyEHR architecture: main modules, databases and their interactions

The Data Management System handles data storage and retrieval from and to the chosen Ehr Database. The module is split in two layers: a service-oriented API for managing the data and a driver interface that supports multiple data back-ends (Fig. 14). The biomedical data, though composed according to a standard, i.e., openEHR, can be expressed in the chosen database in different formats such as for instance Entity-Attribute-Value [36], a good choice for table-based storage systems, XML [37] or document-oriented databases [38]. In PyEHR the choice of the format is delayed and given to the developer, that writes the specific driver interface, which allows him to select the format that best suits the chosen database. Every time a record is inserted the Data Management System forwards it to the Index Service. There its structure get normalized, sorted and compared to the existing ones in the Structures Database and inserted if new. Then the existing or new id of the structure is returned to the Data Management System that stores it along with the encoded record into the Ehr Database.

**Fig. 14.**
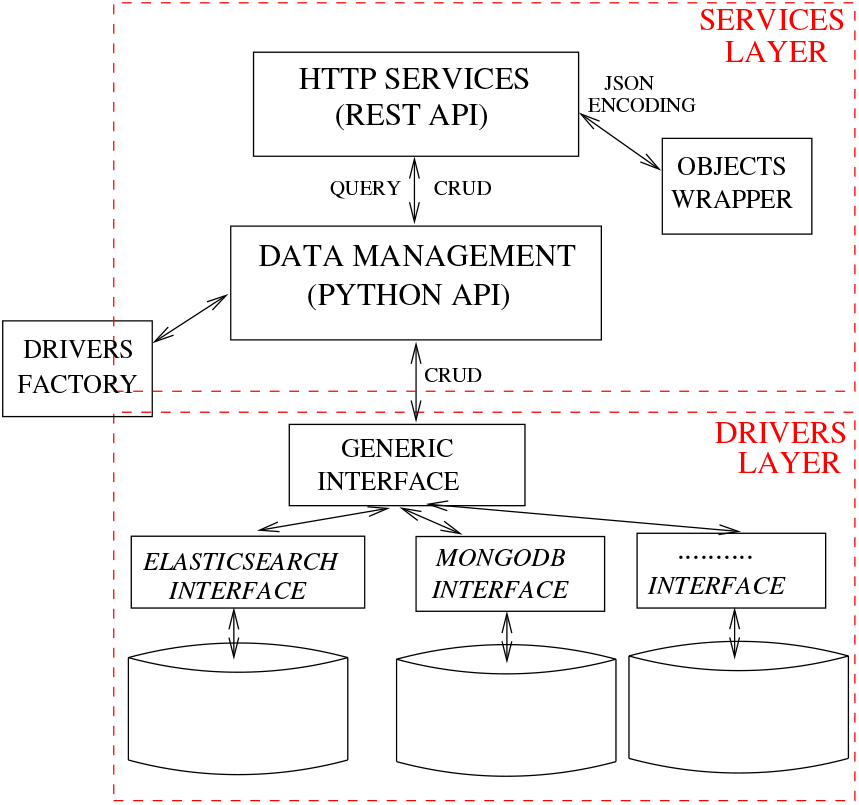
Architecture of the Data Management System.

The Query Management System directs and interacts with different players to get the query results. In figure 15 is shown the path followed by a query and controlled by the Query Manager. The query journey starts at nine o’ clock and follow a clockwise path. The first step involves the input acquisition. The AQL input query is typed by the user in a REST client or submitted, leveraging the python API, through command line interface or a program. The Service Layer of the Data Management System get the query and feed it to the Query Manager Service Core (QMSC) that instantiates a Parser and pass the input query to be processed. The Parser converts the AQL query string into a Query Object Model (QOM) instance that grants access to the query’s data and structure via API calls. The location part of the QOM, that has the CONTAINS clauses, see section VI-B, is then given to the

**Fig. 15.**
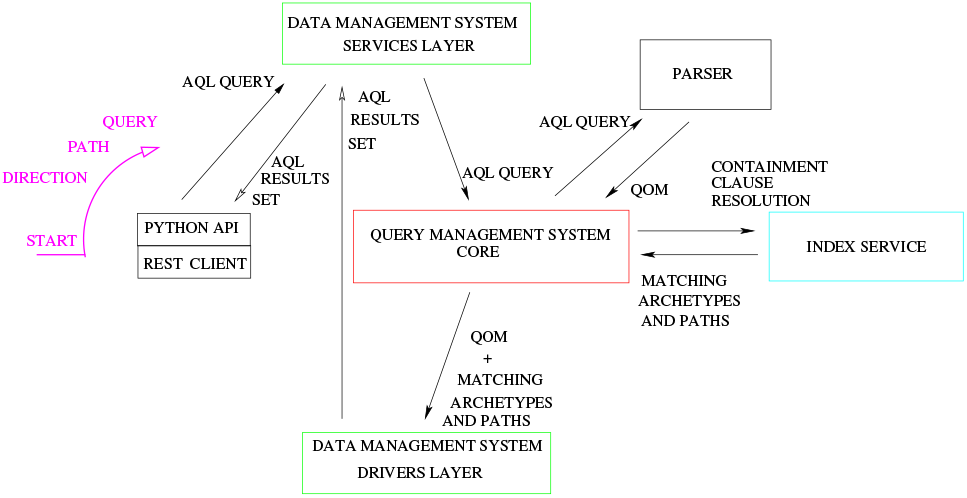
pyEHR query flow

Index Service. A list of matching structures and according paths is returned to the QMSC that sends them together with the QOM to the Driver Layer of the Data Management System. The Driver Layer translates the QOM into the specific query language required by the storage system, executes the query and returns the results as an AQL Results Set, a simple format that represents query results as sets of columns and rows. The results are returned to the user through the Service Layer of the Data Management System.

## VI. Scalability Assessment

### A. Artificial Data Creation

The data creation intended to, above all, generate data that are similar to real ones but possibly more challenging, in order to do a stress test of the system. We want, in other words, to put ourselves in the worst realistic scenario. For the ‘‘constant number of records” test, as said previously, we forged ten millions of EHRs, five millions of which are noise, not responding to any of our queries, and five millions can be seen as five one-million datasets that respond each to their own queries. The reason for the five datasets is to have, along with the single responding dataset needed, four additional responding datasets to check the sensitivity to the random position of the matching records; they answer to the same type of query that yields in all the identical amount of records, which, due to the randomness of the generation, may appear in different places of the structures. For the “constant load” test we chose what we thought was the simplest approach, that is to generate the ten million dataset step by step. We created a ten percent of good data along with ninety percent of noise and replicate the process ten times to have all the needed datasets with the same fraction of records responding to the queries.

Each EHR is written in openEHR formalism, i.e., ADL, and is composed of different archetypes nested to form a tree type structure.

We created complex and varied EHR structures to simulate, through aggregations of archetypes, the output of biomedical pipelines, like for instance that shown for digital pathology in figure 3. The structures have, in json, the appearance of figure 16. Then each structure, drawn randomly from a list of created structures, is instantiated with different values in order to obtain the desired number of matches to the queries. We built, and instantiated, around 2600 unique structures and in figure 17 we show some of them. The circles represent the archetypes and it’s apparent that we can have very deep nested structures. In figure 18 are depicted respectively, from left to right, the maximum relative depth distributions for all the json elements and for the archetypes in our ten millions EHRs along with a normal distribution with the same statistical properties. All the json elements means including both the archetypes and the archetypes’ contained elements so the resulting depth largely increase. From the statistical point of view, we have a mean ‘‘maximum depth” of about 6.7 for the right graph with a standard deviation of 1.7, and a mean of about 66 and a standard deviation of 15.7 for the left one, that is to say that each archetype has roughly a mean maximum depth of 10. The complexity can be high as there are structures that can reach over 110 nested elements and contain up to 12 archetypes in a single branch direction. The maximum depth in the archetype distribution is very similar to a normal distribution.

**Fig. 16.**
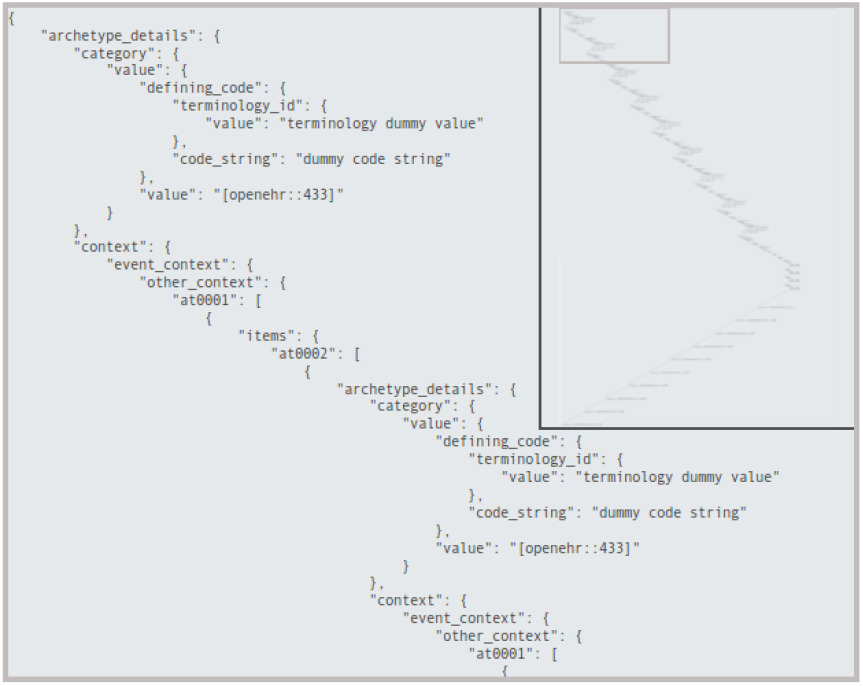
JSON view of a generic EHR structure.

**Fig. 17.**
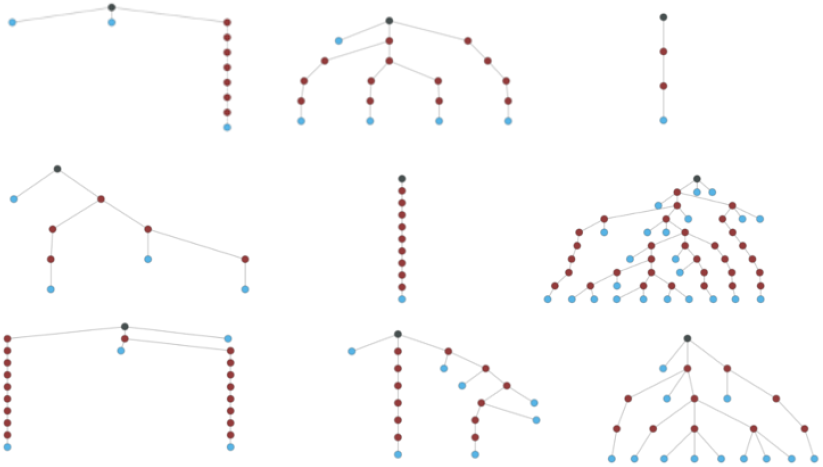
Graph representation of some EHR structures generated.

**Fig. 18.**
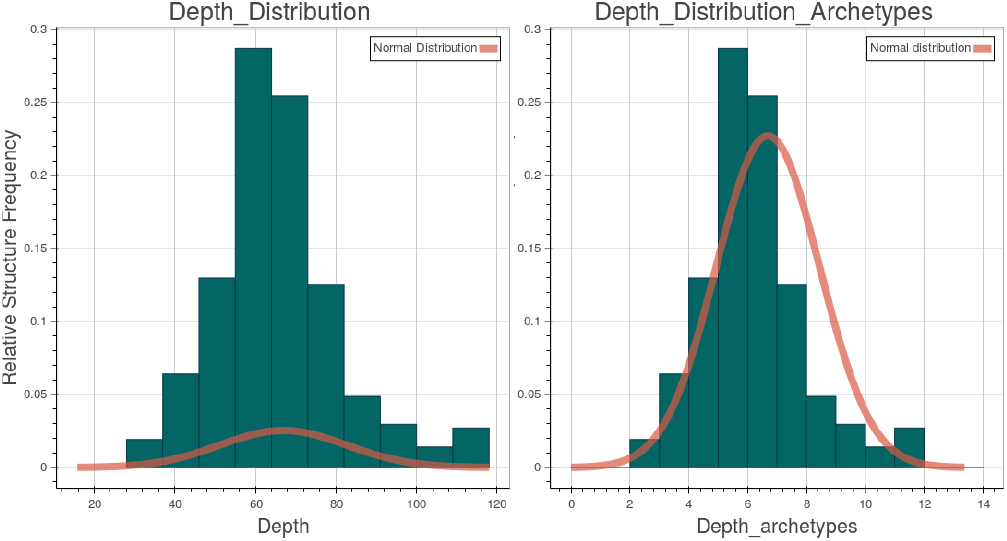
Structures relative depth distribution on the created EHRs.

The maximum width is represented in figure 19. The width is the number of elements measured at each level of the tree structure. The modal value for its maximum is 8 but the width can jump to more than 150.

**Fig. 19.**
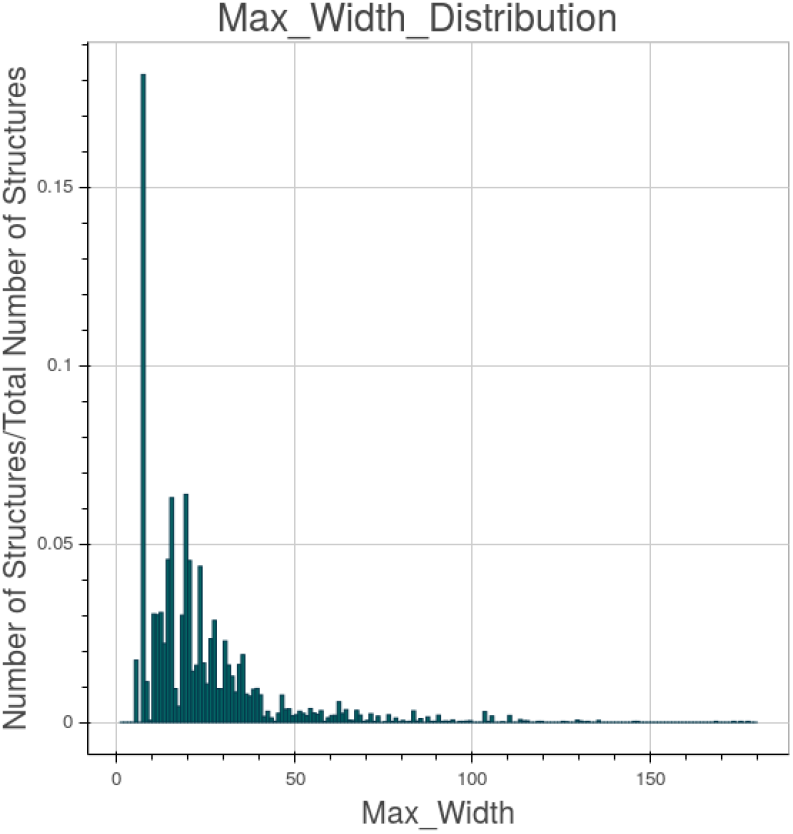
Structures max width distribution on the created EHRs.

### B. Queries Creation

We created five types of AQL queries, to which we assign a number and a brief name for future references, and they are, in ascending complexity: “1-match single”, “2-match multi”, “3-hit single where”, “4-hit multi and where”, “5-hit multi or where”. The first query has a single element in both SELECT and FROM clauses. The second one has two elements to be returned in SELECT clause and a single FROM clause. The third type has a single element in SELECT, FROM and WHERE clauses. The fourth query differs from the third for the existence of a second WHERE clause joined with an AND condition to the first one. The last type has, with respect to the fourth one, the two WHERE clauses in OR. Any FROM clause comes with one or more CONTAINS keyword, meant to add complication to the query and bound to the nested nature of the data structures. As a matter of fact each query is performed on 4 different levels of containment. For instance a level 2 “type 3” query is shown in figure 20. The level can be deducted by the number of CONTAINS keywords. In the example an EHR to match the given query must have anywhere inside its structure the blood pressure observation archetype openEHR-EHR-OBSERVATION.blood_pressure.v1 within a composition archetype called openEHR-EHR-COMPOSITION.encounter.v1.lbl-00001. As already said in section III, between the two archetypes there may be any number of archetypes, included none. Depending on the number of results we decided to let each query yields a simple count of the matching EHRs or both a count and the matching records. The threshold for fetching the results has been set to 10k records, hence under that value the records are counted and returned. The first two types of queries have a number of results that goes from 175k records, at level 5, to 1M records, at level 2, and therefore only a count is performed on them. The last three types range from 1.75k, at level 5, to 10k, at level 2, so both count and fetch are executed. It has to be underlined that the records that match a query at a given level, automatically match the query at any of the upper levels.

**Fig. 20.**
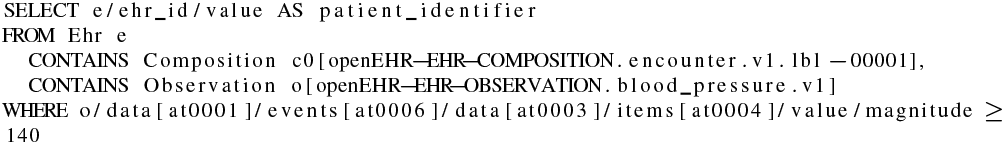
Example of a created level 2 query of “type 3”.

### C. Comparison Reference

A comparison reference is dictated by the need to understand if the measured performances magnitude order is promising. Among the full spectrum of the solutions to our problem we can imagine two diametrically opposite approaches. The first one is a relational database management system RDBMS to store all of our data. The problem is the retention of their relationships. That calls for a very complex arrangement of tables, supposing it achievable, an ad hoc solution with a delicate balance that any modification, addition or removal of data or relationships may easily break. That would involve a complex schema hard to maintain and update. The second approach is classifiable somewhat as a brute force technique. It implies the use of a big data oriented, and therefore easily scalable, software for searching on all the data. We chose the latter approach, that is the safest and simplest one, to have a meaningful comparison and put our results in perspective. As a software we adopted Apache Hadoop Mapreduce [39], one of the most well-known “big data” frameworks for writing scalable applications, already adopted in medical “big data” processing [40]–[43]. The data, put in the hadoop file system, hdfs, are crunched through a recursive search algorithm to find the ones that match the given query. We restricted the comparison to the most simple queries, the “1-match single” queries, with all their different levels of containment.

### D. Data Insertion

The data insertion for the two NoSQL databases, and in particular for Elasticsearch, has proved to be a process very sensitive to parameters tuning. Given that the aim of our work was about the querying performances we did not spend too much time on it but in production it is an area that definitely deserves special attention. The time scale for the insertion of the ten million records, from scratch, are on the order of a couple of days for MongoDB, about four days for Elasticsearch and just the time to write to the disk in hdfs for Apache Hadoop, that is, for the cluster we used, see section VI-F, on the order of about 30 minutes.

### E. Results: Constant Number of Records

The “constant number of records” test, CNR for brevity in the ensuing paragraphs, is, as the name tells, a test where the number of records keeps unchanged while varying the number of working nodes. It has been conducted on Amazon Web Services(AWS). The AWS chosen instances, labeled “r3.2xlarge”, can be placed, for characteristics, in the middle of the family of the so-called “Memory Optimized Instances”. Each node has 8 virtual High Frequency Intel Xeon E5-2670 v2 Ivy Bridge 2.5GHz Processors, 61 GiB of RAM and the storage ensured by one SSD disk of 160 GB.

Twelve machines have been used in the test. Ten machines are the working horses of the software and two are the minds devoted to the control of the operations. In particular within the Apache Hadoop framework one machine is the namenode, responsible for the management of the distributed storage HDFS and the other runs YARN, the Resource Manager that receives and runs application in the cluster depending on the available resources, and the JobHistoryServer, a server that manages past completed/killed jobs. The other ten machines are both datanode and nodemanager, so they store the data and perform the queries. It has to be underlined that ten nodes have always been committed to the storage in Apache Hadoop Mapreduce, for replication necessity, even when we used fewer nodes for the calculations. In PyEHR calculations the two control nodes are used for the main program, i.e., pyehr, and for the database master. The other ten are used as slaves by the database manager system to store, query and retrieve the data. In contrast with Apache Hadoop Mapreduce, the number of nodes that actually store the data can assume any value from one to ten as we add nodes.

We’d like to point out that before choosing the final hardware instance many tests have been carried out within the Apache Hadoop Mapreduce framework. In particular we tried to evaluate the impact of disk, cpu, bandwidth and memory on the performances for the chosen problem to give Apache Hadoop Mapreduce fair hardware conditions to operate. We made tests changing the number of disks (from one to twelve), the type of disks (SSD vs HDD), the bandwidth (1Gbps, 10Gbps), the amount of memory (from 16 up to 121 GB) and the number of processors (from 8 to 20). The net result was that the problem, for Apache Hadoop Mapreduce, is mainly cpu bounded, given the right amount of ram, so we opted for a final AWS instance that was a good compromise between these requirements and the desire to consider only commodity hardware.

Two notes before moving on to the results. The first one is that in all pictures that will follow we represented as a point the mean value of the calculations and as a bar their standard deviation. The number of repetitions for each computation has been set to 10, however, except for the spread curves, all five datasets have been used to enrich the statistics upping the repetitions considered to 50, that is, in other words, 50 values per point/calculation. The second, and final, note is to underline that we are showing, mostly, the count results because the fetch, though needed, it’s only a trivial data upload operation. Moreover the count is more challenging because Apache Hadoop Mapreduce performed it very efficiently in its reducer phase.

#### 1) Apache Hadoop Mapreduce

In figure 21 is shown the behaviour of Apache Hadoop Mapreduce for a “type 1” count query. The results for the four different levels are contained within few seconds in a graph that reach the extreme of 1000 seconds for the single node cluster to drop to about 100 seconds for the ten nodes cluster.

**Fig. 21.**
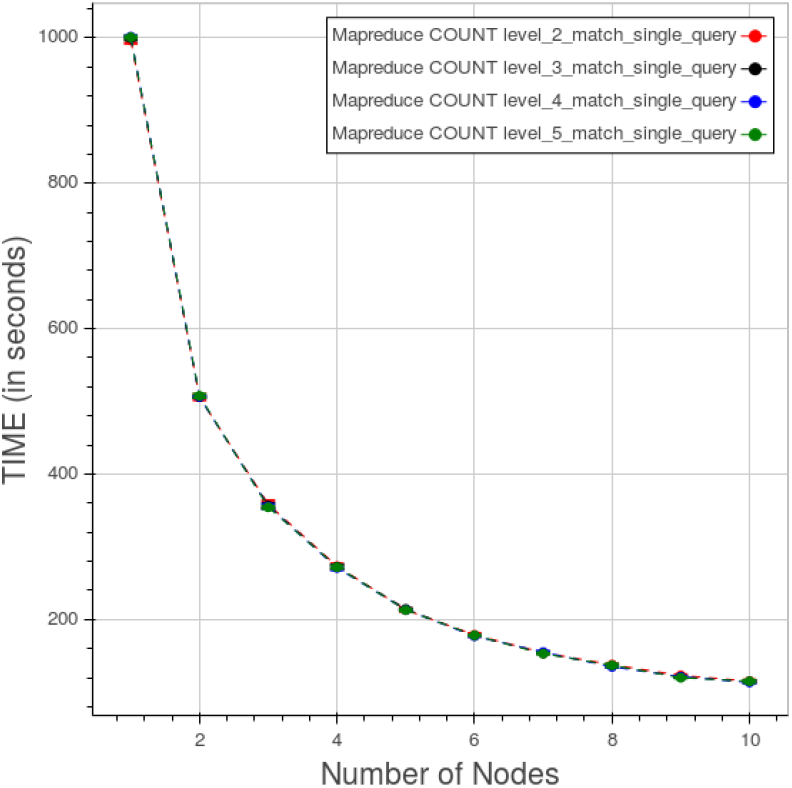
Apache Hadoop Mapreduce. CNR. Time vs number of nodes for a “type 1” count query at different levels.

The spread of the results, due to considering separately the five datasets, at level 5 is displayed in figure 22. The curves are very close with the farthest data, at worst, four seconds apart.

**Fig. 22.**
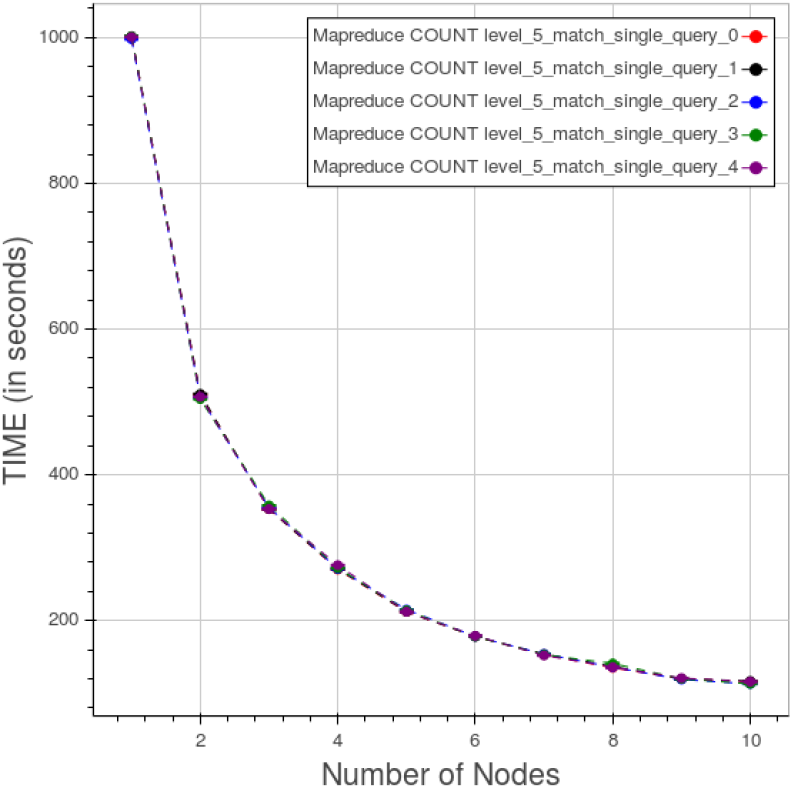
Apache Hadoop Mapreduce. CNR. Data spread for a “type 1” count query at level 5.

#### 2) PyEHR with MongoDB

Figure 23 exhibits the behaviour of PyEHR with the MongoDB driver for a “type 1” count query at the four different levels. The low level queries take much time because they involve a greater number of records.

**Fig. 23.**
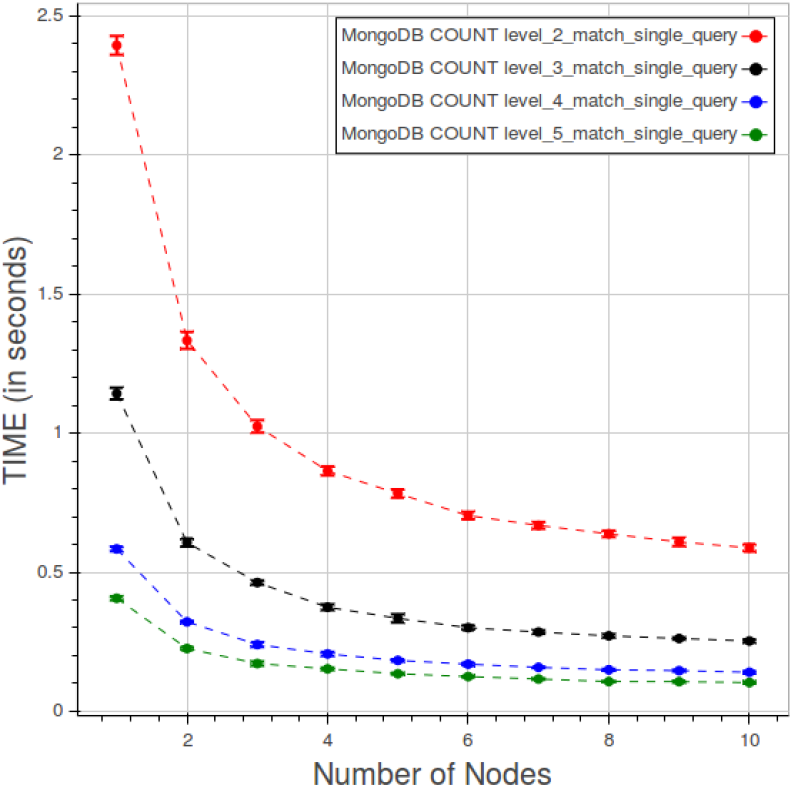
PyEHR with MongoDB. CNR. Time vs number of nodes for a “type 1” count query at different levels.

In figure 24 the different types of count queries are plotted at level 5. The query of “type 5” with two WHERE clauses in OR is the most time consuming while the query of “type 4” with two WHERE in AND and the “type 3” with one WHERE are very close to each other. The queries without WHERE are clearly faster than the rest.

**Fig. 24.**
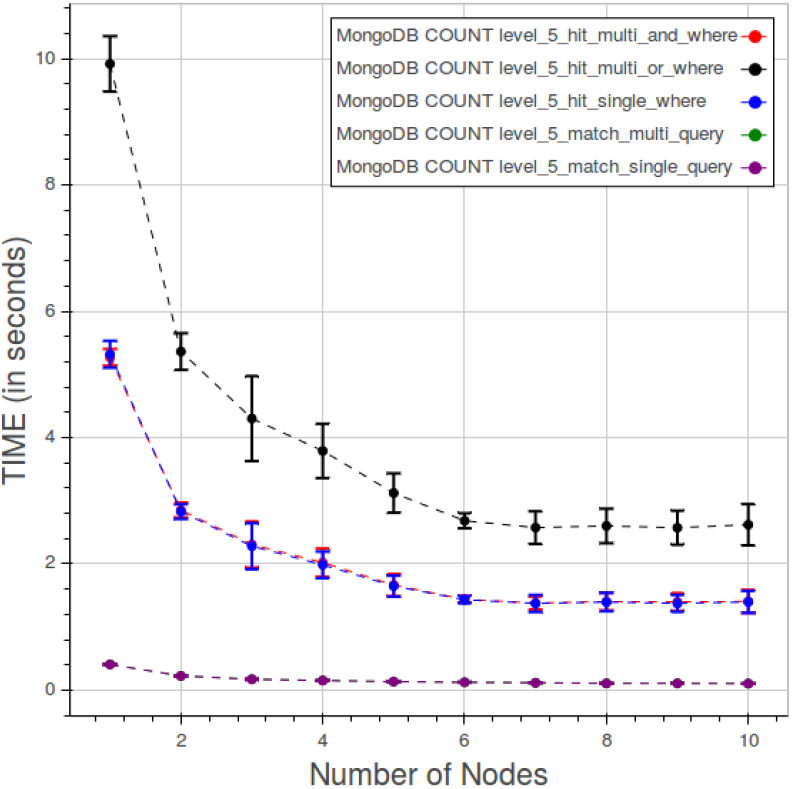
PyEHR with MongoDB. CNR. Time vs number of nodes for different type of count queries at level 5.

Figure 25 indicates how, in a query of “type 3”, the elapsed time is divided between the index, count and fetch phases. The indexing consumes a very little, close to constant, amount of time. Time in the counting and fetching curves improves as we add nodes, rapidly up to five nodes than the curves get almost steady.

**Fig. 25.**
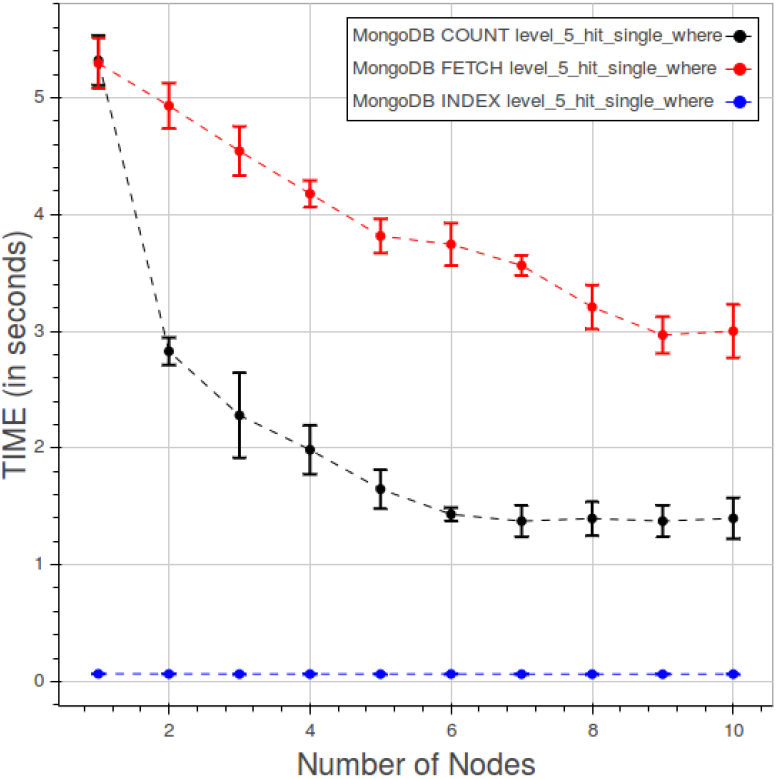
PyEHR with MongoDB. CNR. Time quota of index, count and fetch operations in a “type 3” query at level 5.

The spread of data among the datasets, deducible from figure 26, is rather restrained.

**Fig. 26.**
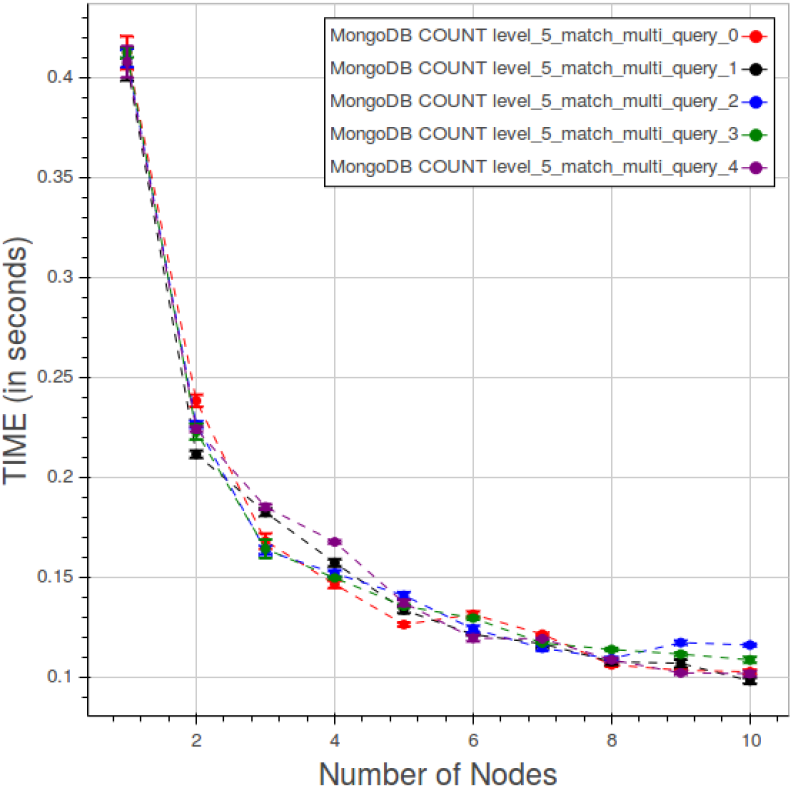
PyEHR with MongoDB. CNR. Data spread for a “type 2” count query at level 5.

#### 3) PyEHR with Elasticsearch

The behavior of PyEHR with the Elasticsearch driver for a “type 1” count query at different levels is displayed in figure 27. The scale of the chart is very small with respect to the other driver and even more compared to the Apache Hadoop Mapreduce graph. However, as in the MongoDB case, time lowers as we go up with the levels. For any given level, the results seem to improve up to a certain point, that depends on the level, and then the curve flattens.

**Fig. 27.**
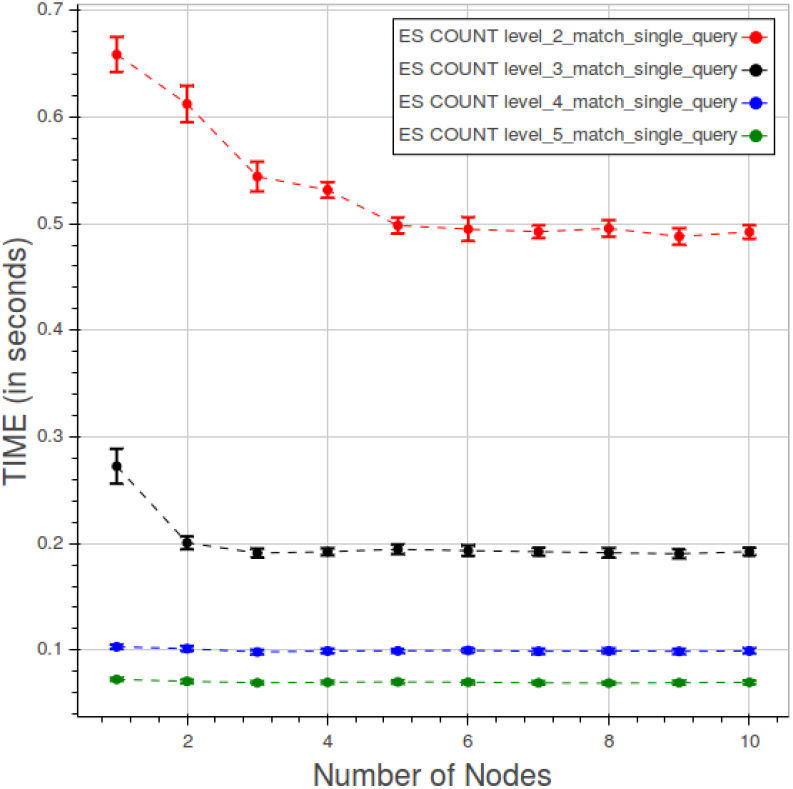
PyEHR with Elasticsearch. CNR. Time vs number of nodes for a “type 1” count query at different levels.

In figure 28 are plotted the curves for the five types of count queries at level 5. The scale is smaller than the previous ones so the variations must be evaluated in that perspective. We can see two cluster of curves, one for “type 1” and “type 2” queries ad the other for the remaining ones. Time quota for indexing, counting and fetching are shown in figure 29. As expected the fetching takes much more time than the other two operations. In this case indexing can’t be neglected, due to the low overall figures.

**Fig. 28.**
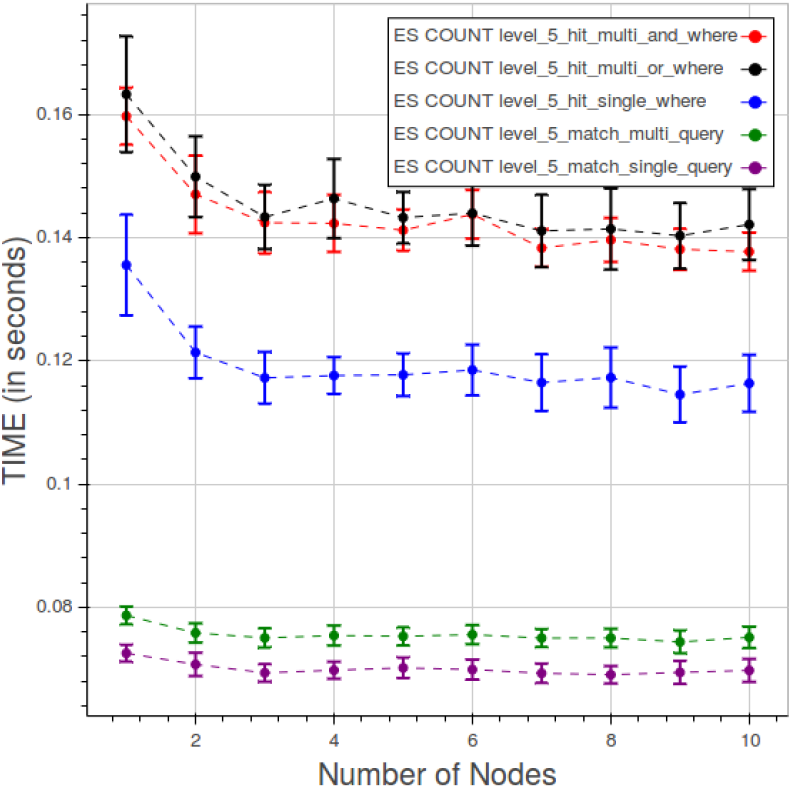
PyEHR with Elasticsearch. CNR. Time vs number of nodes for different type of count queries at level 5.

**Fig. 29.**
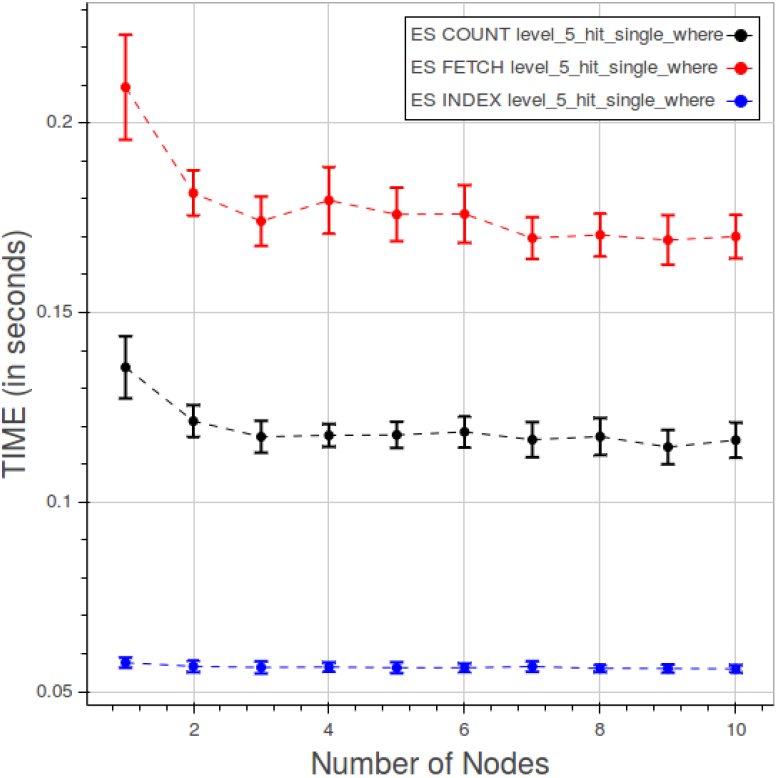
PyEHR with Elasticsearch. CNR. Time quota of index, count and fetch operations in a type 3 query at level 5.

The distribution of points for a “type 2” count query at level 5 is displayed in figure 30. The curves are closely gathered, with a maximum separation between the farthest datasets around 9 thousands of seconds.

**Fig. 30.**
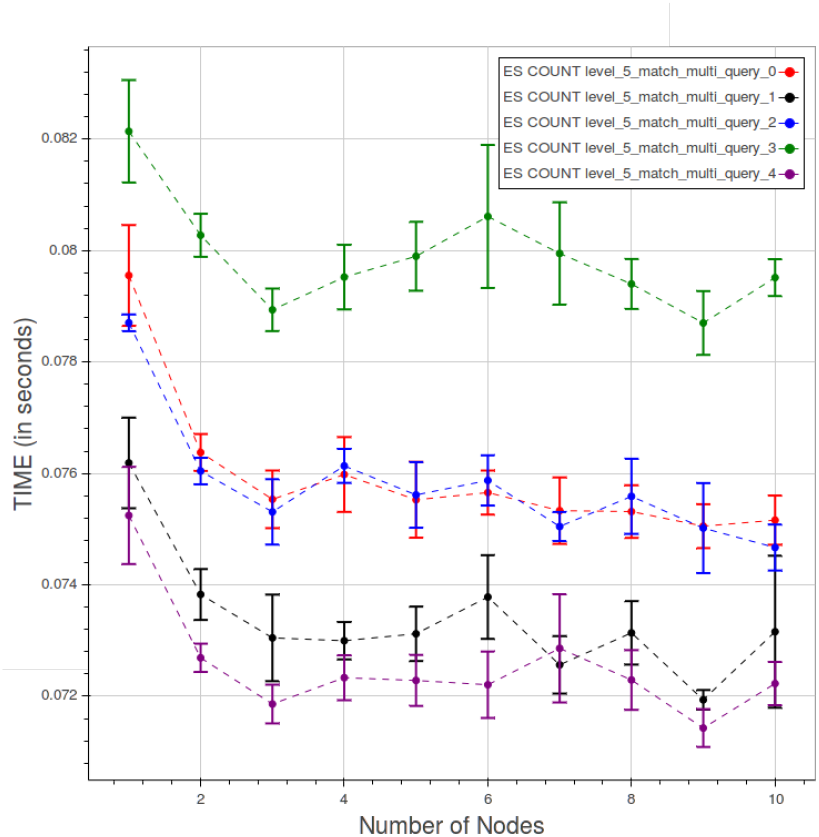
PyEHR with Elasticsearch. CNR. Data spread for the 5 datasets for a “type 2” count query at level 5.

### F. Results: Constant Load

The “constant load” test, CL for conciseness in the following, demands a concurrent linear change in both the number of records and the nodes, maintaining constant the load per machine, hence the name. It has been carried out on a local private cluster of the CRS4 research center. The private cluster has nodes with 8 cpu Intel Xeon E5440 2.83GHz, 16GB of RAM and one HDD of 240GB. Alike the CNR test, twelve machines have been used in the test, ten for storing and doing the actual computations and two for control and queries launching. More details on that role division have been given previosly in VI-E. Five repetitions have been done for each calculation. As before, we chose to put in the graphs mostly the count results.

#### 1) Apache Hadoop Mapreduce

Figure 31 shows the curves for Apache Hadoop Mapreduce for a “type 1” query at different levels. The points are relatively close, apart from the first two nodes where both the distances, between the levels, and the standard deviation, for each level, are large.

**Fig. 31.**
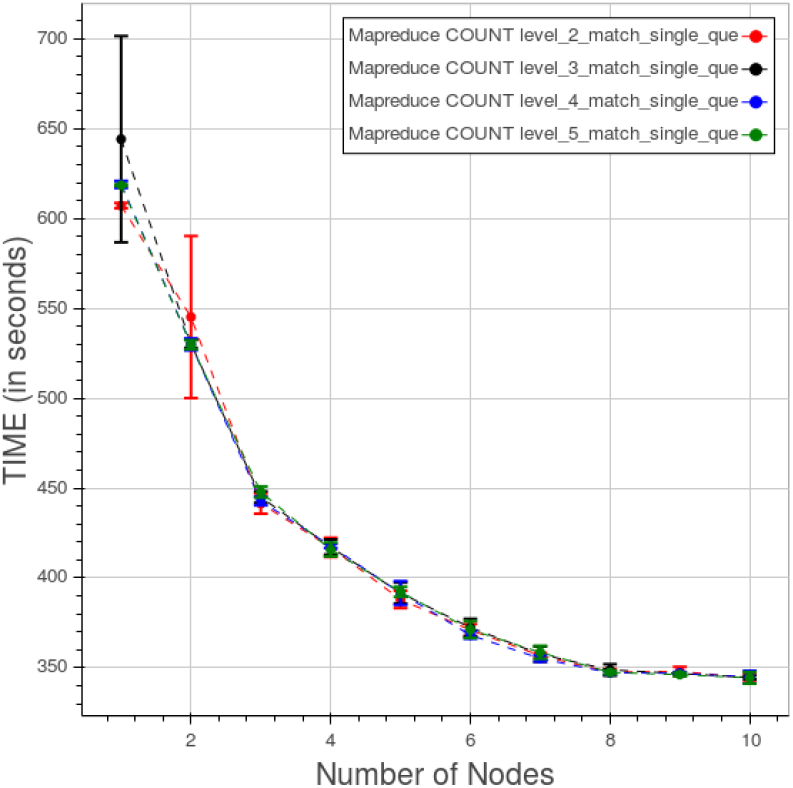
Apache Hadoop Mapreduce. CL. Time for “type 1” count query at different levels.

#### 2) PyEHR with MongoDB

In figure 32 is displayed the behaviour of PyEHR with the MongoDB driver for a “type 1” query at the four different levels. The curves are fully separated.

**Fig. 32.**
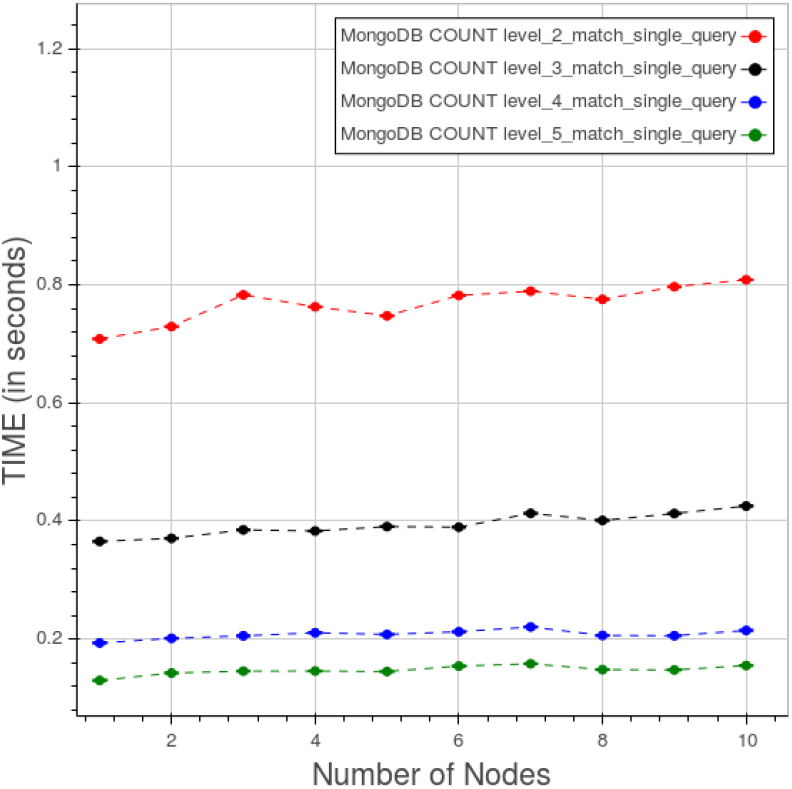
PyEHR with MongoDB. CL. Time for “type 1” count query at different levels.

The effect of changing the type of query at level 5 is shown in figure 33. The “type 3”, with a single WHERE clause, and the “type 4”, with two WHERE clauses in AND, are very close and so are the “type 1”, with a single FROM match, and the “type 2”, with multiple FROM matches. There are anomalies at the node number 5 and 7 that perturb the last three types of query.

**Fig. 33.**
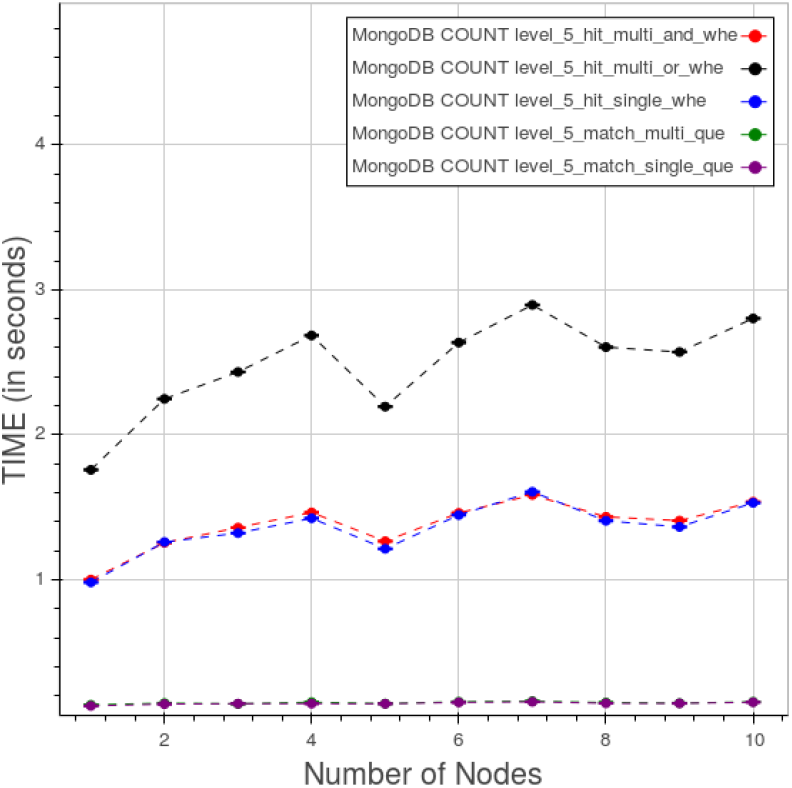
PyEHR with MongoDB. CL. Time for different type of count queries at level 5.

Figure 34 portrays the time consumption ascribed to index, count and fetch phases. While the first two have a steady trend the third one clearly grows with the number of nodes.

**Fig. 34.**
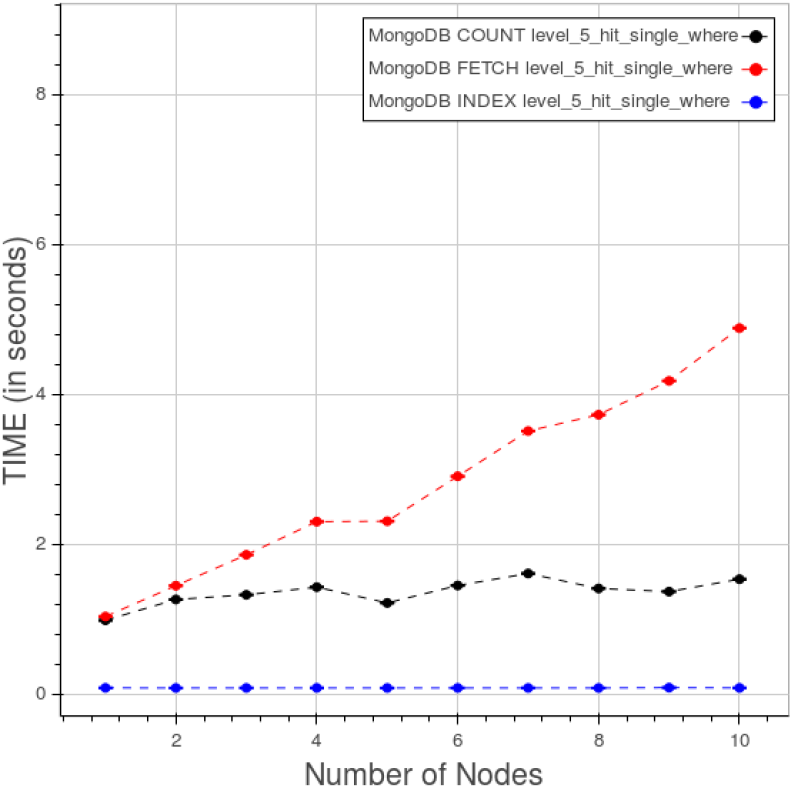
PyEHR with MongoDB. CL. Time quota of count, fetch and index operation in a “type 3” query at level 5.

#### 3) PyEHR with Elasticsearch

The results for PyEHR with the Elasticsearch driver for a “type 1” count query at different levels are presented in figure 35. The curves are neatly separated.

**Fig. 35.**
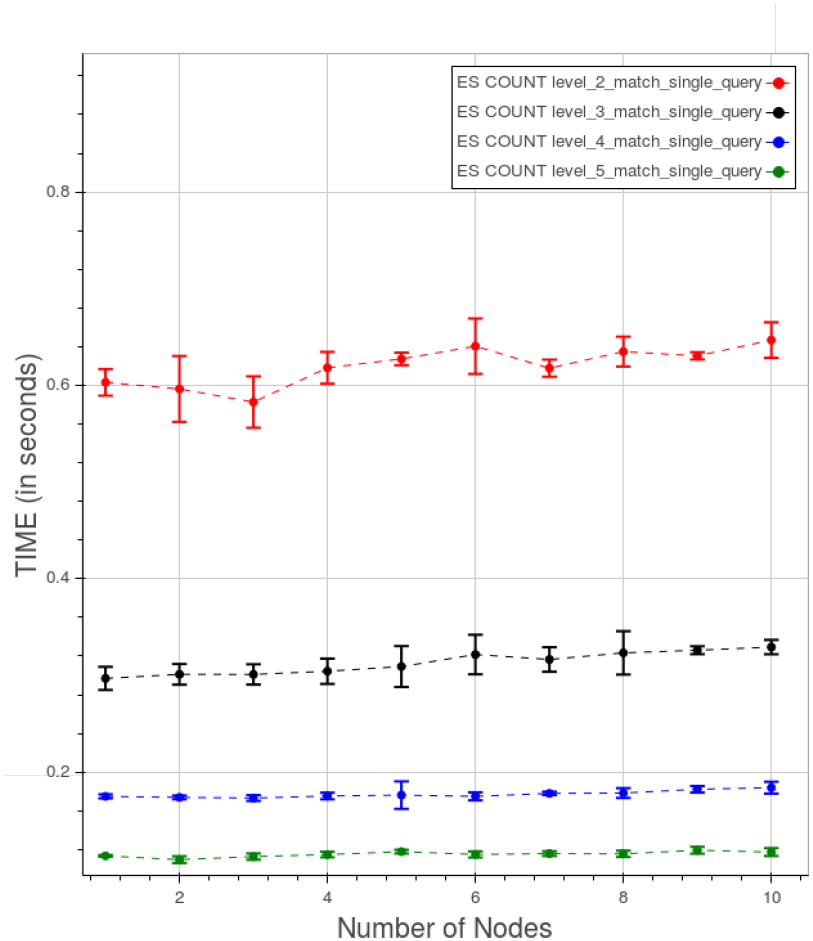
PyEHR with Elasticsearch. CL. Time for type 1 count query at different levels.

In figure 36 is shown the effect on the results of changing the type of query at level 5. The last three types of query clearly are monotonically increasing with the number of nodes, though the figures keep small with respect to both the other driver and Apache Hadoop Mapreduce.

**Fig. 36.**
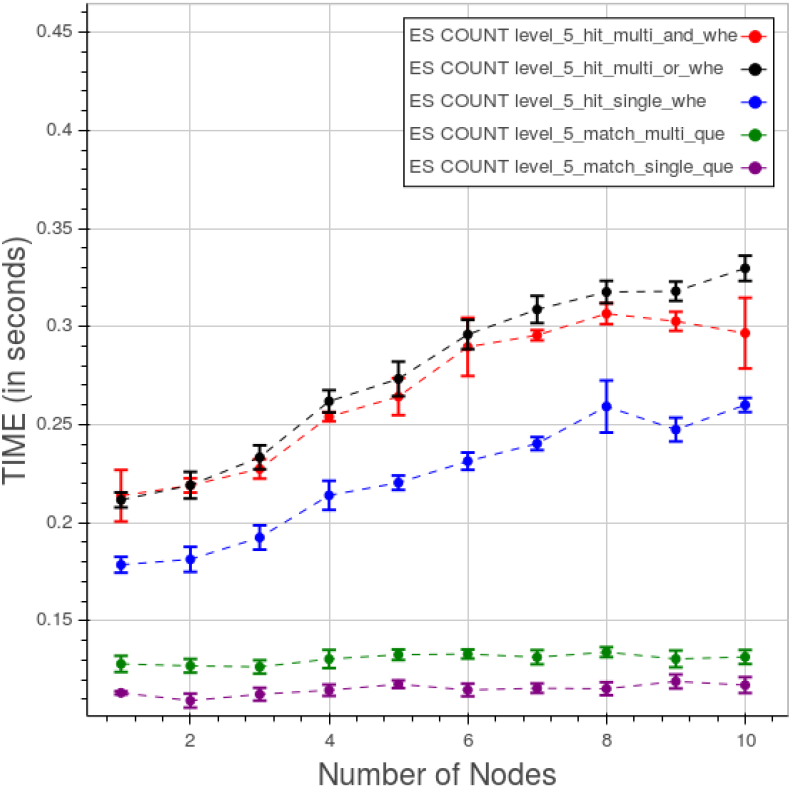
PyEHR with Elasticsearch. CL. Time for different type of count queries at level 5.

Finally figure 37 displays the time quota of index, count and fetch operations for a “type 3” query at level 5. While the indexing keeps a constant trend both the counting and the fetching time increase to some degree as we add nodes and data.

**Fig. 37.**
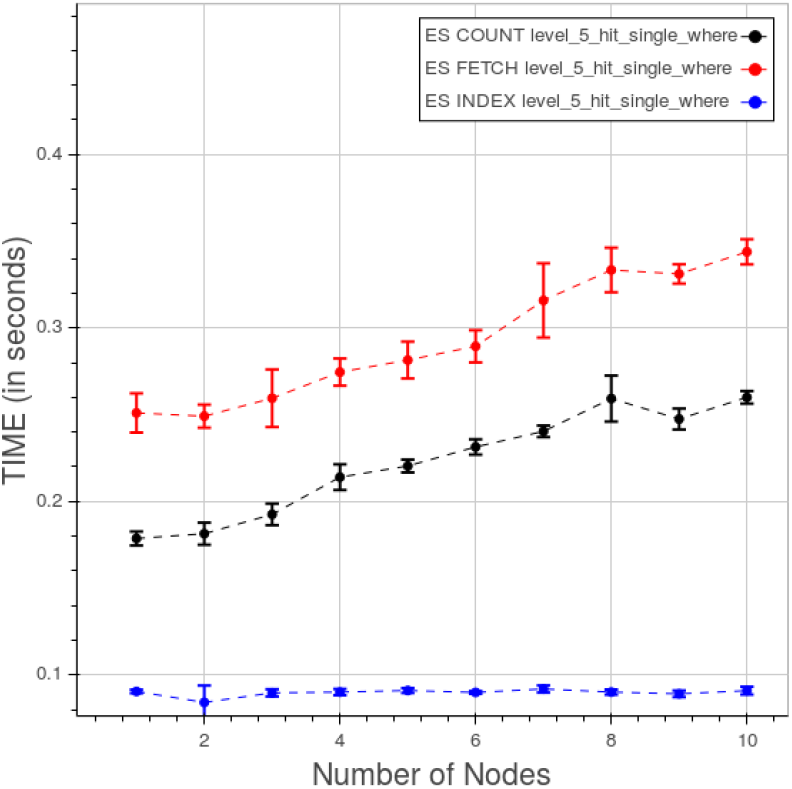
PyEHR with Elasticsearch. CL. Time quota of count, fetch and index operation in a “type 3” query at level 5.

## VII. Comparison between PyEHR and Apache Hadoop Mapreduce and Results Analysis

In figures 38, 39 are shown the comparisons between PyEHR and Apache Hadoop Mapreduce, respectively for the CNR and the CL tests. Figure 38 refers to a CNR type 1 query at level 4. What we expected here was that as long as we add nodes the time had to improve, that is decrease, at least to a certain point. That is the trend for all three curves.

**Fig. 38.**
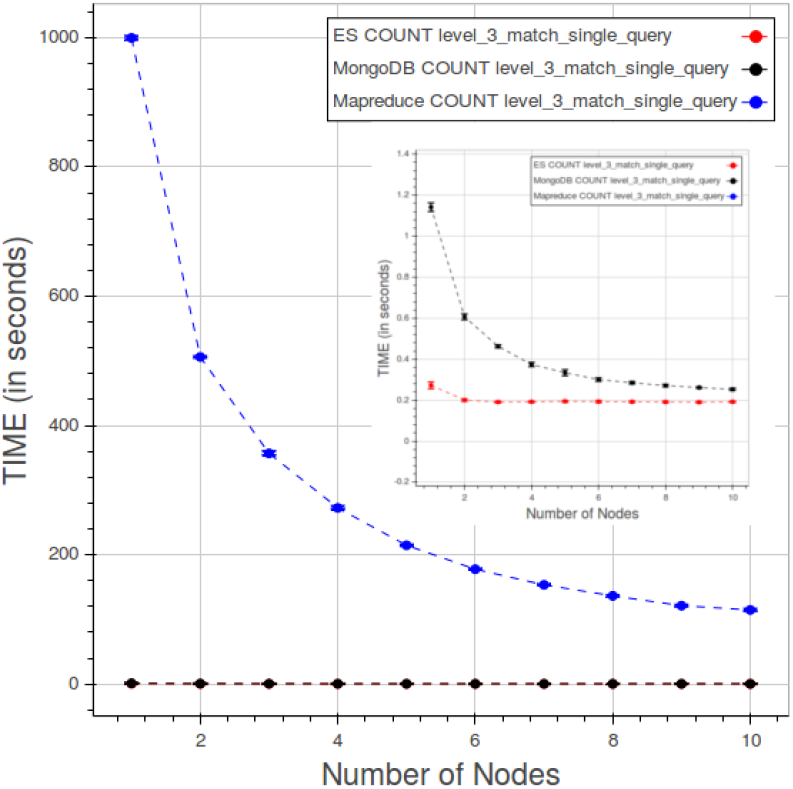
CNR. Results compared for a type 1 query at level 3.

**Fig. 39.**
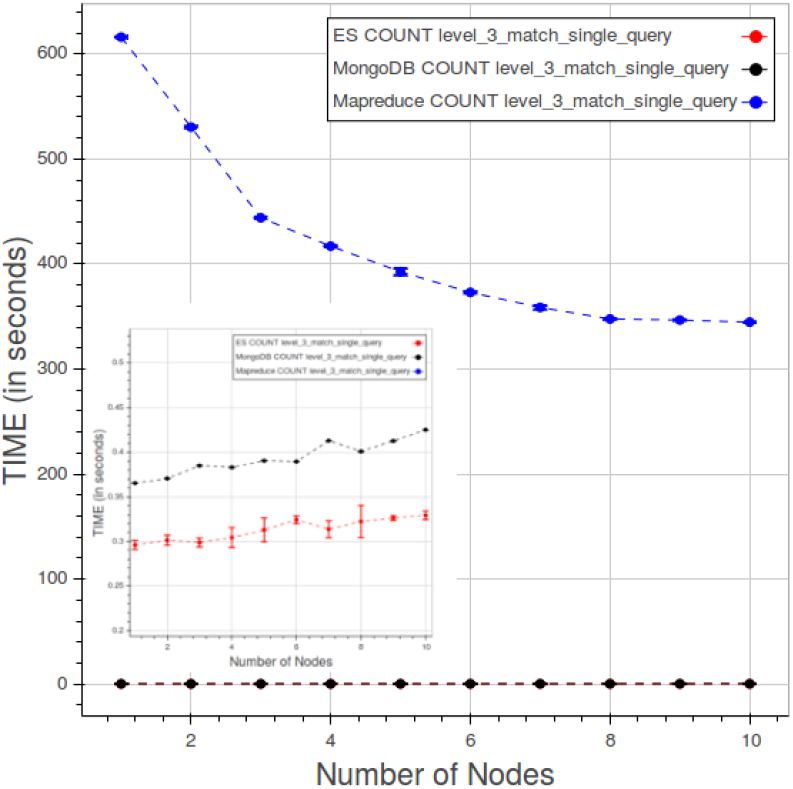
CL. Results compared for a type 1 query at level 3.

By examining the figures 25 and 29 we can assert that the cost of indexing is constant, understandable depending solely on the number of data/structures, whereas the counting and fetching go down as we add nodes, though not indefinitely. The total values for PyEHR, with any driver, and Apache Hadoop Mapreduce are of course very much separated. They differ by two to three orders of magnitude.

The second picture, figure 39 relates to CL test, precisely to a type 1 query at level 3. The desired behavior would be a constant curve where the time value remains the same while adding simultaneously both data and nodes in the same amount endlessly. Apache Hadoop Mapreduce goes even better improving as we add nodes until about eight nodes, then the curve flattens. PyEHR with the two drivers on the other hand gets slightly worse and that is accentuated in the fetch operation, a behavior that can be inferred from the examination of figures 34 and 37. Anyway the numbers are very small and again two to three orders of magnitude smaller than those of Apache Hadoop Mapreduce.

We reckon that the structures indexing done in PyEHR is primarily the reason for this huge magnitude difference between its results and Apache Hadoop Mapreduce ones in both tests. As we have seen in section IV, the query engine receives from the index service a list of the structures that match the given query, each with their path or paths. That allows the driver to filter out the records which possess a structure whose id is not included in the list and therefore to speed up considerably the overall query process. In addition Apache Hadoop Mapreduce has to run through any part of the structures whereas in PyEHR are evaluated only the paths given, for each structure of the list, by the index service, thus saving additional time.

PyEHR seems to behave well with both the drivers. We see a little room for improvement in the parallelization of the index service, though its contribution to the query times in our tests is mostly very small. It has to be reminded, though, that, in nearly absence of configurations tweaking, the time to insert the records is largely in favor of Apache Hadoop Mapreduce.

## VIII. Related Work

Limiting ourselves to non commercial product, there are several pieces of software written to manage clinical data [44]. What is lacking, generally speaking, is applications aimed specifically at managing big amount of complex structured clinical and biomedical data. In particular are missing tools for storing the data preserving their semantics and provenance information and for traversing them horizontally retrieving population information, for secondary use, while most of the software available now performs routinely vertical retrieval of medical records per single patient. Most of the work currently available is based on relational database management systems whose scalability and flexibility are severely put to test by large collections of complex heterogeneous structured data.

Among the openEHR based EHR management projects we would like to mention EHRflex, LiU EEE, ResearchEHR and Ethercis.

EHRflex [45], [46] is an archetype-based clinical registry system designed to employ clinical archetypes as guidelines for the automatic generation of web interfaces, oriented to a clinical use and data introduction. It resorts to eXist, an open source native xml database, to store the data. As stated in Miranda et al Paper [47] XML databases, at least the ones tested in the article, without further optimizations are not suitable as persistence mechanisms for openEHR-based systems in production if population-wide ad hoc querying is needed, being orders of magnitude slower than the relational databases considered.

LiU EEE, described in the paper by Sundvall et al. [48], is an EHR environment designed for educational and research purposes. The product is expressly made to help newcomers and developers experiment with and learn about the openEHR model, from ADL archetype handling to AQL queries. The system, not intended for large-scale analysis, relies on the xml DBMS eXist.

Ethercis or Ethereal Clinical Information System is a system that allows simple interactions with clients using RESTful API and persists data in a separate DB engine, supporting both relational and NoSQL DBMS. Currently it’s implemented the interface for PostgreSQL 9.4. Queries are written in SQL and mixed with json functions and operators.

Other openEHR solutions include yourEHRM [49] by Atos Research that uses MongoDB as internal DBMS and a piece of software cited in paper by Austin et al. [50] that adopts PostgreSQL and it’s the core inside both an academic product named Cortext and a commercial one called HeliconHeart.

Finally it’s worth to be cited the work by Miranda Freire et al [51] where three NoSQL XML databases, BaseX, eXistdb and Berkeley DB XML, and one document oriented NoSQL database, Couchbase, are compared with a relational database, MySQL for health care datasets up to 4.2 million records. Couchbase outperforms, for the bigger dataset, the others on a single node. The distributed configuration is discussed only for Couchbase.

## IX. Conclusions and Future Work

In this paper we have described PyEHR, a data access layer designed to help the creation of clinical and biomedical data management systems for secondary use. Special attention has been put into the design of a system able to cope with large collections of complex heterogeneous structured data, preserve their information and perform on them efficient and scalable queries to get actionable insight. The main features of the system are therefore its support of openEHR standard, that allows to handle very heterogeneous structured data maintaining their provenance and semantic content and interrogate the data at the archetype level, and the indexing of structures, which speeds up the whole queries task. PyEHR’s scalability has been assessed through two kinds of querying test: constant number of records where the records are kept constant while varying the number of nodes and the constant load that sets a fixed number of records per node. Ten million artificial records with complex structures have been generated for each of the tests conducted and five type of queries have been written and tested. The results tell that PyEHR has good scalability properties with both drivers and when compared to a program written for Apache Hadoop Mapreduce, a well-known big data framework, PyEHR outperform it on both tests’ categories.

Future work will be probably focused on upgrading our data access layer to Elasticsearch 2.0 and extending the AQL support. Another foreseen improvement is the parallelization of the index service.

PyEHR is freely available, distributed open source on GitHub at https://github.com/crs4/pyEHR

## X. LIMITATIONS OF THIS WORK

This study does not simulate a real production scenario with concurrent access to the system and insertion, updates and queries intermixed. Essentially in our tests the queries are executed sequentially.

The data are not from real EHRs but, as told in VI-A, we tried hard to make them challenging from the querying point of view. The queries does not include “ORDER BY” and “TIMEWINDOW” clauses as they are not yet implemented.

The acid properties of the system are about the same of the underlying database. We added the external, to the database, versioning support for our EHRs with optional rollback of previous versions.

The results both for the data insertion and querying are the product of little of no configurations tweaking so they are not supposed to be the best results achievable as there’s, for sure, room for improvement.

To assess the scalability we used the following software: Apache Hadoop 2.6.0, MongoDB 3.04, Elasticsearch 1.5.0, BaseX 8.0.1.

With respect to the openEHR implementation, in order to get a fully compliant system there should be a data and query validation service that fetches archetypes/template from a local/remote repository, a terminology translator and optionally a data templates provider.

## Acknowledgments

One of the authors, Giovanni Delussu, has performed his activity in the framework of the International PhD in Innovation Science and Technology at the University of Cagliari, Italy.

